# Protein phosphorylation and oxidative protein modification mediate plant photosystem II disassembly and repair

**DOI:** 10.1101/2023.05.03.538416

**Authors:** Steven D. McKenzie, Sujith Puthiyaveetil

**Affiliations:** Department of Biochemistry and Center for Plant Biology, Purdue University, West Lafayette, IN 47907, USA

**Keywords:** PSII repair cycle, thylakoid protein phosphorylation, STN8, PBCP, protein oxidative modification

## Abstract

The light-driven water-splitting reaction of photosystem II exposes its key reaction center core protein subunits to irreversible oxidative photodamage. A rapid repair cycle replaces photodamaged core subunits in plants, but how the large antenna-core supercomplex structures of plant photosystem II disassemble for repair is not currently understood. Phosphorylation of reaction center core protein subunits has been suggested as a mechanism of disassembly. Consistent with this, we find specific involvement of phosphorylation in removing peripheral antenna from the core and monomerization of the dimeric cores in *Arabidopsis*. However, photosystem disassembly occurred to some degree even in the absence of phosphorylation as suggestive of other unknown mechanisms of disassembly. Here we show that the oxidative modifications of amino acid residues in core protein subunits of photosystem II are active mediators of disassembly. Exogenously-applied hydrogen peroxide induces photosystem disassembly, especially the conversion of the monomeric cores into two reaction center subcomplexes. We further show that the extent of monomer disassembly is proportional to the oxidative protein damage, with the fully disassembled reaction center subcomplexes containing more modifications. In the monomeric core, some amino acid oxidative modifications map at the D1-CP43 interface as consistent with a dissociation of the core along these subunits. Oxidative modifications thus likely disassemble only the damaged monomeric cores, ensuring an economical photosystem disassembly process. Our results suggest oxidative protein modification represents an ancient mechanism of photosystem disassembly, and that phosphorylation originated later in evolution to impart explicit control over the repair process.

## Introduction

Oxygenic photosynthesis harnesses the potential of two nearly inexhaustible raw materials – sunlight and water – to produce readily usable chemical energy in the form of ATP and NADPH. Critical to light energy conversion in oxygenic photosynthesis is the ability of photosystem II (PSII) to extract electrons and protons through a light-driven water-oxidation reaction. The electrons derived from water are used by PSII to reduce the electron carrier plastoquinone (PQ) to plastoquinol (PQH_2_), and the protons contribute to the proton motive force that drives ATP synthesis. The molecular oxygen, a byproduct of water-oxidation, acts as a terminal electron acceptor in aerobic respiration to release the chemical energy contained in photosynthates. A Mn_4_CaO_5_-cluster is key to PSII’s remarkable water-splitting chemistry. However, in the absence of electron sinks and sources, as under intense sunlight or abiotic stresses, incomplete water oxidation and charge recombination events in PSII produce highly reactive hydrogen peroxide (H_2_O_2_), hydroxyl radical (HO^•^), and superoxide anion (O_2_ ^•−^) (1, 2). The HO^•^ and O_2_ ^•−^ oxidatively modify a number key amino acid residues in the D1 and D2 reaction center proteins of PSII, impairing PSII function and causing photoinhibition – the light-induced loss of photosynthetic activity (1, 3–6). Early irreversible oxidative modifications of amino acid ligands of the Mn_4_CaO_5_-cluster and the nonheme iron trigger a cascade of additional modifications in D1 and D2, as consistent with photoinhibition arising from photooxidative damage at both donor and acceptor sides of PSII (1).

A robust repair process replaces the photodamaged D1 reaction center protein with a newly synthesized copy through an intricate cycle of disassembly of the holocomplex, degradation of the damaged D1, *de novo* synthesis, processing, and insertion of new D1, and reassembly of the holocomplex (7–10). In plants, the repair of photodamaged D1 protein presents two major challenges (11). One of them is the disassembly of the large 1.4 MDa supercomplex of PSII, formed by an extensive light harvesting complex II (LHCII)-based peripheral antenna and a dimeric reaction center core (12, 13). The monomeric plant reaction center core comprises 20 protein subunits, with the D1 protein found buried at the core of the complex (13, 14). The second challenge for repair of plant PSII is to mobilize the photodamaged supercomplexes from tightly stacked granal thylakoid regions to unstacked stroma lamellae for degradation of photodamaged D1 and co-translational assembly of nascent D1. Disassembly of the holocomplex might solve both of these problems as disassembled smaller complexes are presumably more mobile (15, 16). Though many aspects of plant PSII repair cycle have been well understood, little is known about the mechanism of disassembly. Two serine/threonine protein kinases State Transition 7 (STN7) and 8 (STN8), respectively, phosphorylate PSII in a light quality and quantity dependent manner (17–21). Though these two kinases act with some redundancy, STN7 predominately phosphorylates the peripheral trimeric LHCII protein in the acclimatory response of state transitions while STN8’s major targets are the PSII core protein subunits (17–19). PSII phosphorylation is reversible as a PSII Core Phosphatase (PBCP) dephosphorylates core phosphoproteins while a separate phosphatase targets the phospho-LHCII (22–24). Core protein phosphorylation by STN8 has been suggested as a mechanism of disassembly but how precisely phosphorylation leads to PSII disassembly remains unknown (25, 26). A phosphorylation map shows strategic location of PSII core phosphosites at the peripheral antenna attachment sites and at the monomer-monomer interface as consistent with a role in removing the peripheral antenna from the core and in core monomerization (27). An analysis of a knockout mutant of STN8, with significantly reduced core phosphorylation, and a double mutant of STN7 and STN8, lacking PSII phosphorylation altogether, surprisingly revealed that phosphorylation accounts for only a fraction of PSII disassembly, as indicative of other unknown disassembly mechanisms at play (19, 25). Using biochemical, biophysical, and mass spectrometric approaches in model plant *Arabidopsis thaliana*, here we show how core protein phosphorylation acts synergistically with protein oxidative modification for the sequential and economical disassembly of PSII during its repair cycle.

## Results

### Phosphorylation drives peripheral antenna dissociation and core monomerization

To examine the precise role of core phosphorylation in PSII disassembly, we first determined the core phosphorylation level in *Arabidopsis thaliana* wild type (WT), a double knockout mutant of STN7 and STN8 (*stn7stn8*), a knockout mutant of STN8 (*stn8*-*1*), an overexpressor of STN8 (*STN8oe*), and two knockout mutants of PBCP (*pbcp*-*1* and *pbcp*-*2*) under growth light (∼150 μmol m^-2^ s^-1^). Fig. 1*A* shows a phosphothreonine immunoblot of thylakoid samples from these plants. Under growth light, *stn7stn8* shows no core or LHCII phosphorylation as consistent with STN kinases being the sole drivers of PSII phosphorylation (20). *stn8-1* shows a significantly reduced PSII core phosphorylation and a slightly lower LHCII phosphorylation, in line with earlier observations of STN8’s major kinase activity towards PSII core proteins and a minor but physiologically relevant phosphorylation of LHCII (19, 28). The residual core phosphorylation in *stn8-1* (Fig. 1*A*) is likely to be the result of STN7 activity, given the slight substrate overlap and redundancy between the two PSII kinases (20, 21). *STN8oe* and *pbcp* knockout mutants have hyperphosphorylated cores as expected (22, 29, 30).

**Figure 1.**
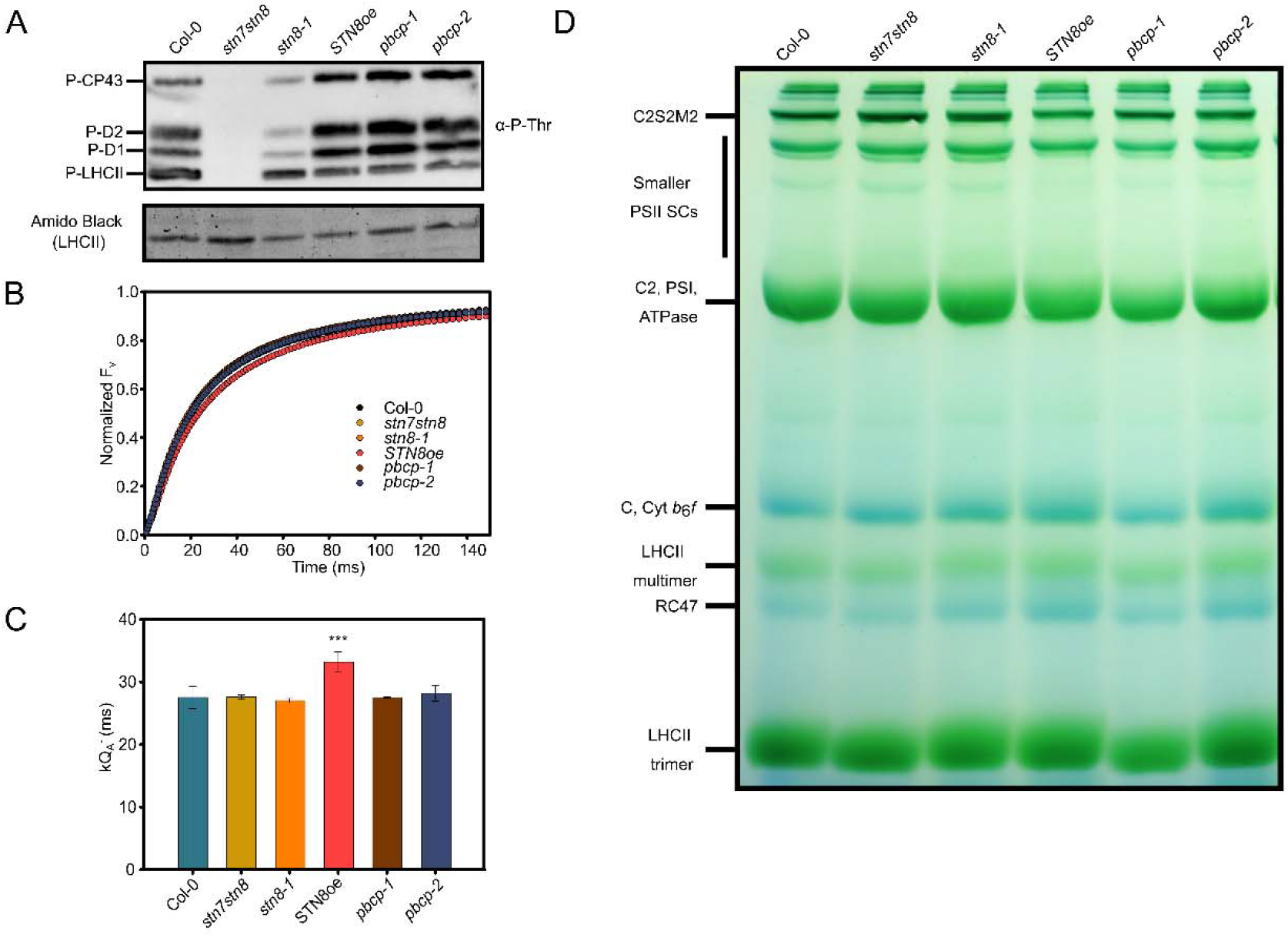
PSII antenna size and assembly status as a function of core phosphorylation level in growth light. (*A*) A phosphothreonine immunoblot of thylakoid samples from wild type (Col-0) and mutant plants. Bands corresponding to four major PSII phosphoproteins are indicated on the left. The amido black staining of the corresponding PVDF membrane is shown below the blot as a loading control with the LHCII protein band labeled. (*B*) Normalized variable chlorophyll florescence yield (F_v_) is plotted against time for wild type and mutant thylakoid samples. Each fluorescence trace is an average of three biological replicates with each biological replicate measured in technical triplicates. (*C*) The rate constant of Q_A_ reduction (kQ_A_^-^) in wild type and mutant thylakoids. Error bars represent standard deviation of three biological replicates (n = 3). Triple asterisks (***) denote a statistical significance (p-value) of <0.001 vs. the wildtype control as determined by a one-way ANOVA test followed by a post-hoc Holm-Sidak t-test. (*D*) A representative blue native gel of wild type and mutant thylakoid samples. Bands corresponding to major thylakoid protein complexes are indicated on the left.

After determining the basal PSII phosphorylation level of all genotypes under growth light (Fig. 1*A*), we then analyzed their PSII antenna size using chlorophyll fluorescence induction (Fig. 1*B*). The room temperature chlorophyll fluorescence arises from chlorophyll a molecules associated with PSII antenna. The rise of the fluorescence from a minimal (F_o_) to a miximal (F_m_) value in the presence of the electron transport inhibitor DCMU (3-(3,4-dichlorophenyl)-1,1-dimethylurea) indicates the progressive reduction of the primary quinone electron acceptor Q_A_. Under a uniform illumination by a green actinic light, a faster rise in chlorophyll fluorescence indicates better excitonic connectivity between PSII units owing to a larger peripheral antenna (31). Fig. 1*B* shows normalized chlorophyll fluorescence induction of isolated thylakoid samples. It is apparent that the *STN8oe* has a smaller PSII antenna size, consistent with its hyperphosphorylated cores causing the accumulation of smaller disassembled PSII. The other mutants, in contrast, show no significant changes in chlorophyll fluorescence induction under the growth light condition (Fig. 1*B*). Fig. 1*C* shows the rate constant of Q_A_ reduction (kQ_A_^-^), which is the inverse time required to reach 60% of the fluorescence rise (31). With its smaller and less connected antenna, *STN8oe* has the highest kQ_A_ ^-^ value while other mutants show no statistically significant difference with wild type (Fig. 1C). It is noteworthy that the *pbcp* mutants, containing hyperphosphorylated cores, show no discernable decrease in antenna size (Fig. 1B and C) in agreement with an earlier study (30). Though the reason for a smaller antenna in *STN8oe* but not in *pbcp* is not clear, it is possible that an overexpression of STN8 results in core phosphorylation and disassembly of a larger percentage of PSII supercomplexes than in *pbcp* mutants, which has a STN8 activity that is similar to wild type.

To account for the possibility that chlorophyll fluorescence induction analysis is not sensitive enough to detect subtle changes in PSII functional assembly, we analyzed the abundance of various disassembly intermediates of PSII by blue native polyacrylamide gel electrophoresis (BN-PAGE). Thylakoid membranes were first isolated from plants grown under growth light and the thylakoid protein complexes were solubilized with the mild detergent n-Dodecyl α-D-maltoside (α-DM, Anatrace). The solubilized protein complexes were subsequently resolved on a blue native gel. For PSII, at least four structural species can be discerned on the gel (Fig. 1D) (32, 33). The largest structural species comprises a group of antenna-core supercomplexes, migrating as multiple bands on top of the gel. The most intense band is a PSII supercomplex known as C2S2M2, consisting of a dimeric reaction center core (C2), two strongly bound trimeric LHCII (S2), and two moderately bound trimeric LHCII (M2). The S-trimers are tethered to the core through the monomeric LHCII protein CP26 while the M-trimers associate with the core via monomeric CP29 and CP24 LHCII proteins. C2S2M2 is the predominant assembly form of plant PSII in low light condition (34). A dimer of C2S2M2 migrates above the C2S2M2 band, and the bands below correspond to smaller PSII antenna-core supercomplexes with a decreasing complement of the peripheral antenna (32, 33). The other three structural species of PSII are a dimeric core, a monomeric core, and a sub-monomeric complex called CP43-free core or Reaction Center 47 (RC47). RC47 contains D1, D2, and CP47 core subunits but is devoid of the CP43 reaction center protein (8, 35, 36). Other thylakoid protein complexes are also resolved on the gel and so are a multimeric and trimeric LHCII complexes (Fig. 1D). It is apparent from the gel that *stn7stn8* and *stn8-1* mutants have slightly more C2S2M2 species than wild type while the *STN8oe* has markedly lower abundance of this supercomplex. The *pbcp* mutants have slightly lower level of C2S2M2 as the wild type. The smaller PSII antenna-core supercomplexes are also less abundant in *STN8oe* and *pbcp* mutants, compared to wild type (Fig. 1D). Another major change in PSII disassembly in *STN8oe* and *pbcp* mutants is a lower proportion of dimeric cores with a corresponding increase in monomeric cores. The abundance of the RC47 complex is higher in *STN8oe* and in *pbcp* mutants.

Since photodamage and core phosphorylation correlate with light intensity (37, 38), we checked whether the disassembly differences between the genotypes become more apparent under a higher light intensity. We thus measured core phosphorylation level and abundance of PSII disassembly intermediates in plants treated with 800 μmol m^-2^ s^-1^ white light for two hours. This moderate light-treated plants show lower LHCII phosphorylation and STN7-dependent core phosphorylation, as expected from an inhibition of STN7 kinase activity at higher light intensities (Fig. S1) (39). STN8-mediated core phosphorylation persists under high light with *STN8oe* and *pbcp* knockout mutants showing hyperphosphorylated cores. However, the extent of differences between the genotypes in PSII antenna size and abundance of different disassembly intermediates were more or less similar to that observed under growth light (Fig. S1).

The steady state level of PSII structural species in vivo is a function of both disassembly and reassembly processes (8). To examine the effect of phosphorylation on disassembly alone, we studied PSII abundance in isolated *in vitro*-phosphorylated wild type thylakoids as they lack PBCP and a few other repair components (22). Wild type plants were first treated with far-red light for 2 hours at 21 °C, followed by immediate isolation of thylakoid membranes from these plants. The far-red light, by its preferential excitation of PSI, causes an overoxidation of the PQ pool and thereby a deactivation of STN kinases (20, 40). One aliquot of far-red-treated thylakoids was incubated at 21 °C with 25 μM ATP under 150 μmol m^-2^ s^-1^ white light, a second aliquot was treated with the same light and temperature conditions but with no ATP present, and a third aliquot kept at dark was used as control for far-red light treatment. Samples were withdrawn from the white light-treated +/-ATP thylakoid samples at 15, 30, and 60 minutes of incubation and analyzed for PSII phosphorylation and abundance. Fig. 2*A* shows a phosphothreonine immunoblot of the far-red light control, white light plus ATP, and white light minus ATP thylakoids.

**Figure 2.**
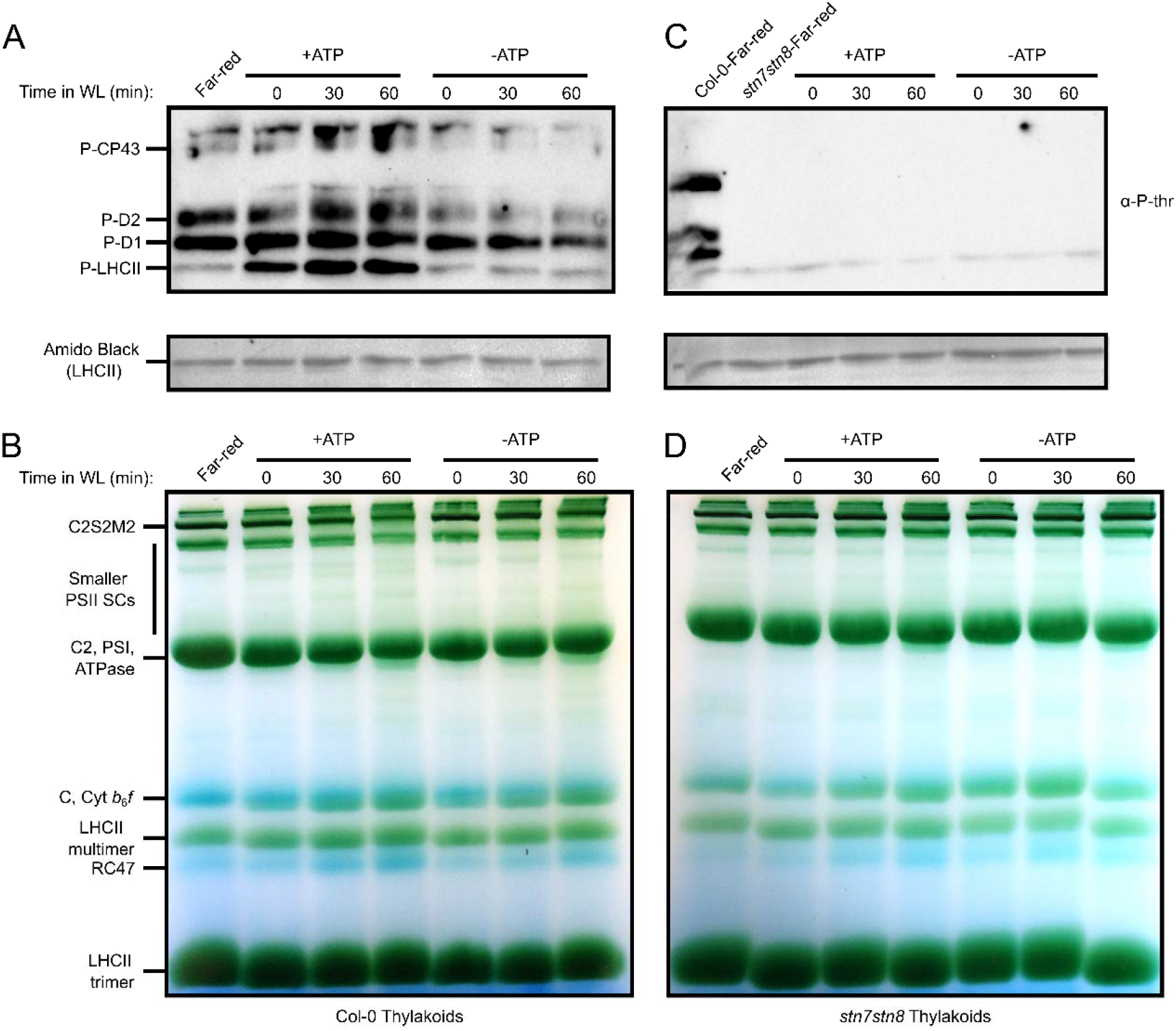
In vitro phosphorylation of thylakoids leads to PSII disassembly. A phosphothreonine immunoblot of wild type (*A*) and *stn7stn8* (*C*) thylakoids incubated with or without ATP in light. Major PSII phosphoproteins are indicated on the left. The corresponding PVDF membrane has been stained with amido black and given below the blot as a loading control. A representative blue native gel of wild type (*B*) and *stn7stn8* (*D*) thylakoids treated with or without ATP in light. Major thylakoid protein complexes resolved are indicated on the left. Time (in minutes) under white light (WL) illumination is also indicated on top of the gels.

As expected, incubation of the far-red-treated thylakoids with ATP under white light leads to increasing levels of LHCII, D1, D2, and CP43 phosphorylation, compared to far-red control. The time-dependent increase in phosphorylation is especially evident for LHCII, D1, and D2 proteins (Fig. 2*A*). The minus ATP sample apparently shows a time-dependent decrease in PSII phosphorylation when compared with the far-red control. It is possible that this is due to dephosphorylation reactions dominating under ATP depletion. A blue native gel analysis of these thylakoid samples reveals clear differences in the abundance of PSII structural species (Fig. 2*B*). Induction of PSII phosphorylation in plus ATP samples leads to the disassembly of the larger C2S2M2 as a function of incubation time, compared to the far-red control. This is accompanied by an accumulation of smaller PSII complexes, including C1 and RC47 (Fig. 2*B*). In minus ATP samples, the abundance of disassembled complexes was higher than far-red control but lower than plus ATP samples, as consistent with a disassembly mechanism that depends not only on phosphorylation but also other light-dependent processes. As a negative control, the experiment was repeated with *stn7stn8* thylakoids. As expected, thylakoids from *stn7stn8* were not phosphorylated prior to or upon incubation with ATP (Fig. 2*C*). BN-PAGE analysis of *stn7stn8* thylakoids revealed no discernable changes in PSII supercomplex abundance, however, the RC47 complex accumulates in both plus and minus ATP samples upon exposure to light (Fig 2*D*). Together, these results indicate a largely phosphorylation-dependent mechanism for the disassembly of the peripheral antenna from the core and for core monomerization and a phosphorylation-independent mechanism for the further disassembly of the monomeric core.

### Exogenous reactive oxygen species induce PSII disassembly

Since phosphorylation seems to account for only a proportion of PSII disassembly (Figs. 1 and 2), we sought the possibility of whether photodamage, in the form of oxidative protein modification, drives PSII disassembly. It has been recognized that H_2_O_2_, generated by inadvertent redox reactions at the donor and acceptor sides of PSII, is a major source of highly damaging O2^•-^ and HO^•^ radicals (1). We therefore incubated thylakoids from wild type plants with increasing concentration of exogenous H_2_O_2_ at 21 °C for 30 minutes in dark. We then analyzed the abundance of PSII structural species in these thylakoids by BN-PAGE (Fig. 3*A*). The peroxide-treated thylakoids show a notable shift in the composition of PSII, with larger antenna-core supercomplexes being converted to smaller antenna-core supercomplexes for every 10-fold increase in H_2_O_2_ concentration. The most remarkable change under H_2_O_2_ however seems to be the disassembly of the monomeric core into RC47, as judged from the marked changes in band intensities of these complexes (Fig. 3*A*). Since some blue native gel bands contain co-migrating complexes, band intensity changes cannot always be equated with the changing abundance of just one complex. This is particularly true for the core monomer, which co-migrates with the cyt *b*_6_*f* complex. We therefore carried out a D1 immunoblot of the blue native gel for detecting changes in just PSII. The blue native gel was destained and the protein complexes blotted onto a PVDF membrane and subsequently probed with a D1 antibody. The D1 blot (Fig. 3*B*) indeed confirms a peroxide-driven dissociation of the monomeric core into RC47 as suggested by the blue native gel (Fig. 3*A*).

**Figure 3.**
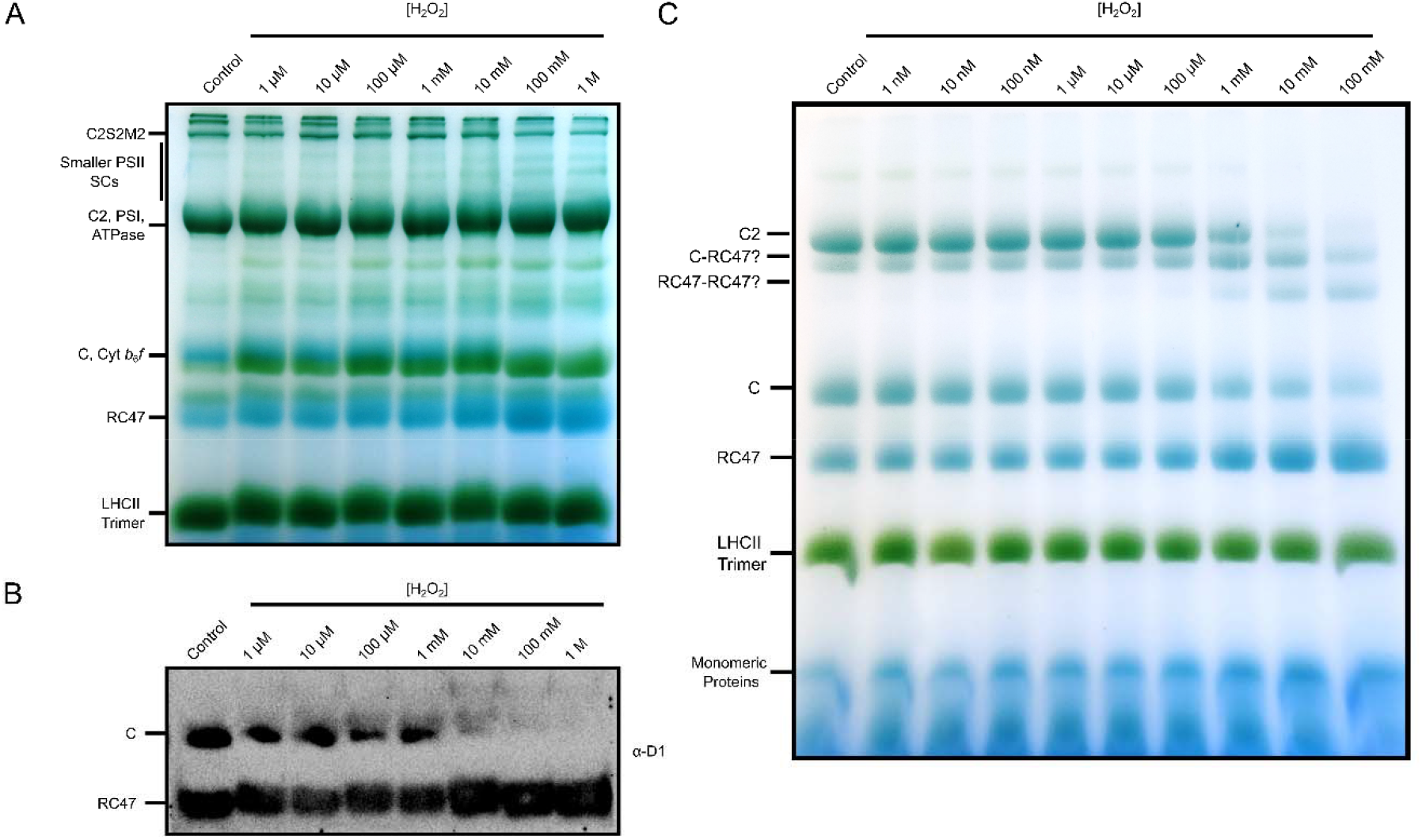
Exogenous H_2_O_2_ treatment results in preferential disassembly of monomeric cores into RC47. (*A*) A representative blue native gel of peroxide-treated wild type thylakoids. Major PSII structural species are indicated on the left. (*B*) A corresponding D1 immunoblot of the blue native gel. The immunoblot has been cropped to show only the C and RC47 bands and the uncropped blot is given in Fig. S2. (*C*) A representative blue native gel of peroxide-treated isolated reaction center cores of *stn7stn8* plants. Disassembly products and copurifying contaminant complexes are indicated on the left. Positions of putative dimeric C-RC47 and RC47-RC47 complexes are also indicated. Smaller complexes or monomeric proteins at the bottom of the gel are likely disassembly products or copurifying contaminants but their identity is uncertain and are labeled as such.

To further examine the specific effect of the peroxide treatment on PSII reaction center core disassembly and to rule out disassembly from other causes such as core phosphorylation and lipid peroxidation, we analyzed peroxide-driven disassembly of isolated reaction center cores from *stn7stn8* double mutant. For this analysis, PSII-enriched granal regions were first isolated from *stn7stn8* thylakoids and solubilized with β-DM. PSII complexes were subsequently separated by anion exchange chromatography and the fraction corresponding to the reaction center core was harvested and concentrated. The purified core fraction was buffer exchanged and treated with different concentrations of H_2_O_2_ at 21 °C for 30 minutes in dark. The peroxide-treated core samples were resolved on a blue native gel (Fig. 3*C*). The core fraction from *stn7stn8* contained both dimeric and monomeric reaction center complexes. Addition of H_2_O_2_ leads to the fragmentation of both dimeric and monomeric cores into smaller subcomplexes as apparent from the gradual decrease in C2 and C band intensities and a corresponding accumulation of RC47 at increasing peroxide concentration (Fig. 3*C*). Remarkably, most of the *stn7stn8* dimeric cores seem to dissociate directly into RC47 rather than first disassembling into a monomeric core intermediate. The new blue native gel bands, found between the dimeric and monomeric cores, may thus represent intermediate disassembly species such as a monomeric core with an RC47 and a dimer of RC47 complexes (Fig. 3*C*). This observation not only reveals how the *stn7stn8* mutant, which completely lacks PSII phosphorylation, disassembles its PSII but also the distinct roles of core phosphorylation and oxidative modification in core monomerization and disassembly of the monomeric core, respectively. The trimeric LHCII appears to be a copurifying contaminant with the core (Fig. 3*C*).

### Innate and peroxide-induced oxidative protein modifications in PSII disassembly

To further elucidate the role of oxidative protein modification in PSII disassembly, we mapped amino acid residues that are oxidatively modified under light (innate) and exogenous peroxide by mass spectrometry. Thylakoids were first isolated from wild type plants grown under growth light and treated with 20 mM H_2_O_2_ or buffer and incubated for 1 hour in dark at 21 °C. The peroxide and mock-treated thylakoids were then solubilized with α-DM and the thylakoid protein complexes separated on a non-oxidizing blue native gel (Fig. 4*A*). The changes in the abundance of C1 and RC47 upon peroxide treatment confirm the earlier observations of peroxide-driven C1 disassembly from regular blue native gels (Fig. 3). Protein bands corresponding to C1 and RC47 complexes were excised from the gel and analyzed by LC-MS/MS. We performed searches for several commonly identified oxidative post-translational modifications in core subunits common to both complexes. Mono-oxidation of amino acid side chain (addition of an oxygen atom) appears to be the most common oxidative protein modification (Tables S1 and S2). Fig. 4B shows the number of unique and shared oxidative protein modifications in C1 and RC47 complexes under H_2_O_2_ and mock treatments as a Venn diagram. Under peroxide treatment, core subunits of RC47 contains substantially more oxidative modifications than those of C1. This observation supports the notion that the increased C1 disassembly seen under peroxide treatment (Fig. 3) proceeds from an accumulation of peroxide-induced oxidative modifications in C1. Under mock treatment, RC47 carries more common modifications than C1. These common modifications are likely induced by the growth light prior to the peroxide or mock treatment of thylakoids and they may account for the smaller extent of C1 disassembly observed in untreated sample (Fig. 3). It is very likely that some of the C1 and RC47 complexes represent reassembly species and may therefore not contain any oxidative modifications. Although the oxidative modifications that are critical for C1 disassembly remains to be identified, the data in Fig. 4*B* suggests two possibilities. C1 disassembly may result from a small set of amino acid modifications that greatly destabilize the monomeric core or a larger set of modifications with each having an incremental destabilizing effect. It is also possible that some of the modifications are unrelated to disassembly (Fig. 4B).

**Figure 4.**
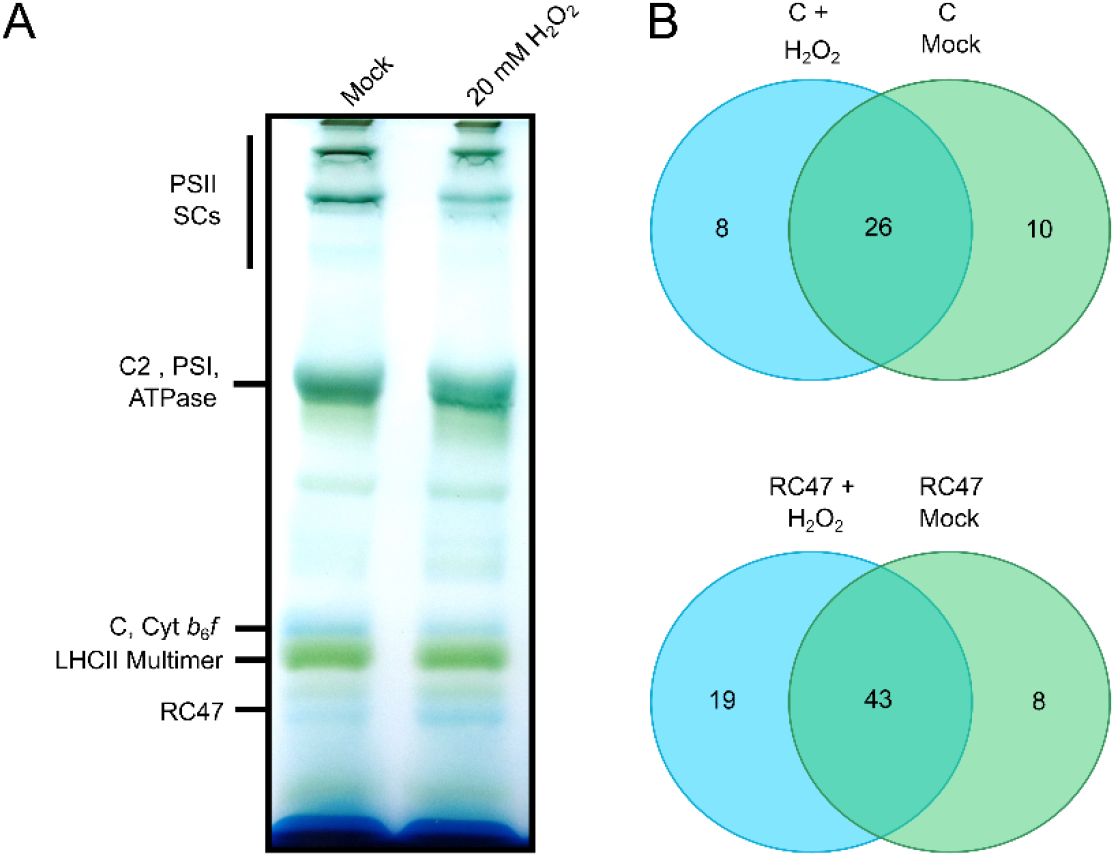
Extent of oxidative protein modification correlates with the degree of PSII disassembly. (*A*) A representative non-oxidizing blue native gel of peroxide-treated wild type thylakoids. Major PSII structural species are indicated on the left. The gel has been cropped to show only one biological replicate. The uncropped gel containing all three biological replicates is given as Fig. S3. (*B*) A Venn diagram showing unique and shared oxidative protein modifications in C and RC47 under peroxide- and mock treatments. The modifications are listed in Tables S2 and S3.

## Discussion

Since its first demonstration nearly four decades ago, the functional implication of protein phosphorylation for PSII structure and function has eagerly been sought (41). Phosphorylation of core protein subunits D1, D2, CP43, and PsbH has generally been referred to as core phosphorylation and the phosphorylation of the peripheral antenna proteins, broadly as LHCII or antenna phosphorylation. Phosphorylation of LHCII is now firmly established as the basis of plant state transitions (20, 21, 40). However, the precise function of core phosphorylation has remained elusive. Various roles have been proposed over the years but the suggestion of an involvement in disassembly is supported by some recent studies (25–27). How exactly phosphorylation leads to PSII disassembly has however remained unclear. Our observations of the overaccumulation of PSII antenna-core supercomplexes in STN knockout mutants and underaccumulation in *STN8oe* (Fig. 1D) confirm earlier findings from other laboratories and are in agreement with the role of phosphorylation in peripheral antenna dissociation as predicted by a phosphorylation map of PSII (Fig. 5A) (19, 25, 27, 29). The increased monomerization of PSII in *STN8oe* (Fig. 1D) is also in line with an earlier study of STN8 (29) and supports the suggestion that core phosphorylation leads to monomerization of the dimeric cores. The slower reduction of Q_A_ in *STN8oe*, the lower abundance of larger PSII structural species in *pbcp* mutants, and PSII disassembly in in vitro-phosphorylated thylakoids (Figs. 1B-D and 2) further strengthen the role of core phosphorylation as a modulator of PSII functional and biosynthetic assembly. It is interesting to ask whether the currently known four core phosphoproteins are sufficient to explain PSII disassembly or are there additional phosphoproteins involved in PSII disassembly. We therefore undertook a phosphoproteomic analysis of wild type thylakoid samples through a phosphopeptide enrichment strategy. Our analysis confirms many of the previously known PSII phosphoproteins and phosphosites (Table S3). However, we also find some novel PSII core phosphoproteins that include PsbL, PsbF, PsbQ-2, and PsbR. The PsbL phosphorylation is especially interesting as this subunit is found at the monomer-monomer interface and is required for core dimerization (42). PsbL phosphorylation may thus drive core monomerization in concert with the two known PsbH phosphosites (Fig. 5A). Since D2 and PsbF phosphosites occur near the putative docking site of PsbS (Fig. 5A), a dissociation of PsbS and or M-trimer may require these phosphorylation events. The function of PsbQ-2 and PsbR phosphorylation is unclear. PsbQ-2 is one of the two isoforms of the lumenal PsbQ subunit of the Oxygen Evolving Complex (OEC) of PSII (43). PsbR is a transmembrane protein that interacts with the OEC protein PsbP (44, 45), and is required for OEC assembly (46). An intriguing possibility is that PsbQ-2 and PsbR phosphorylation is important for the disassembly of OEC from the reaction center core. Indeed, the disassembly/reassembly of OEC is a poorly understood aspect of the repair cycle (8).

**Figure 5.**
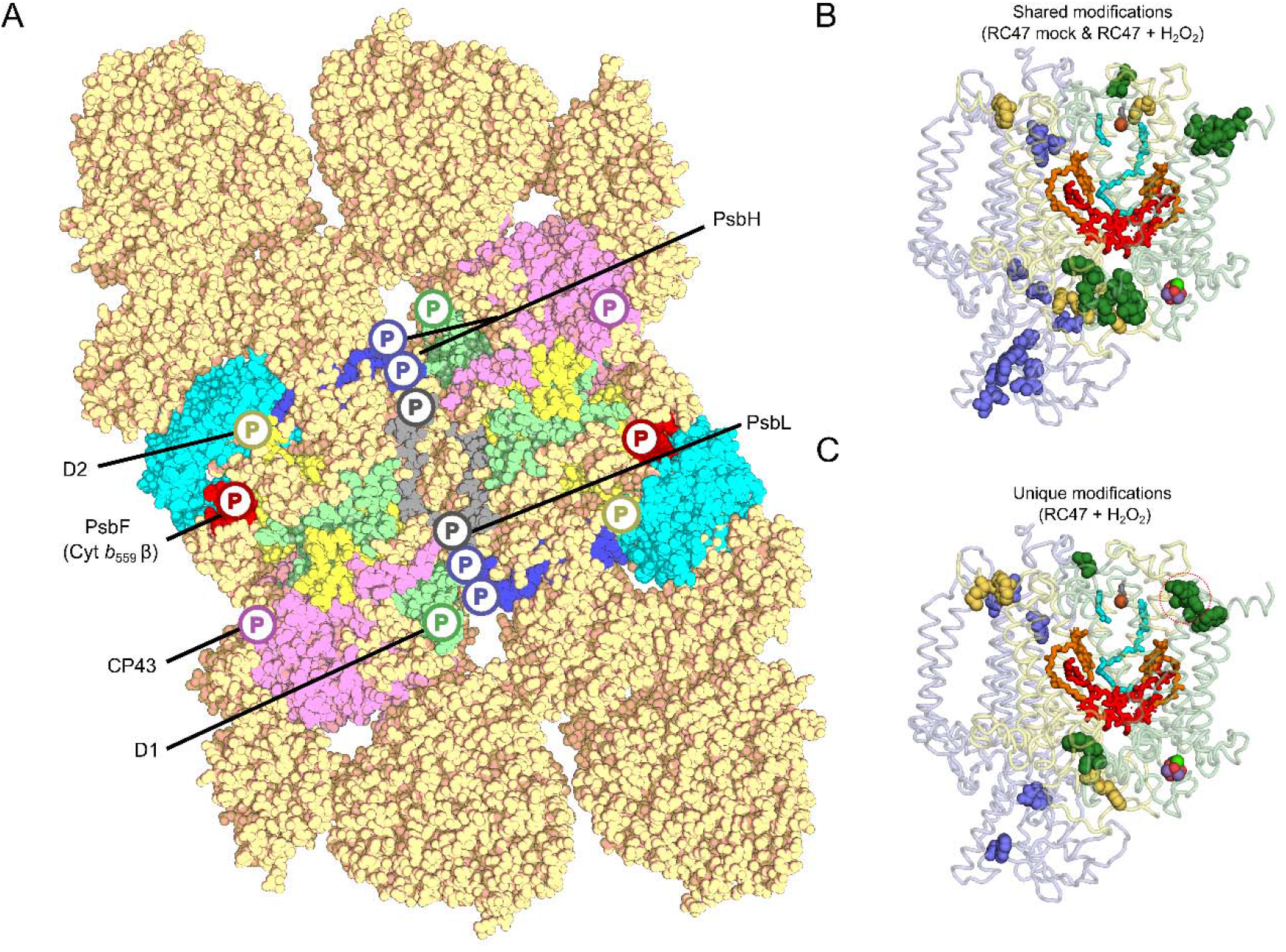
Protein phosphorylation and protein oxidative modification contribute to PSII disassembly. (*A*) A projection of known phosphosites onto plant C2S2M2 supercomplex. The PsbS subunit (in cyan) is also shown according to current predictions on its location (12, 59). Phosphoproteins are rendered in different colors and labeled. D1 and CP43 phosphosites are predicted to drive the dissociation of the S-trimer from the core; and the D2, PsbF, and CP29 phosphosites, the M-trimer. The two phosphosites of PsbH and a phosphosite of PsbL occur at the monomer-monomer interface and they may therefore contribute to the monomerization of the dimeric reaction center cores. The phosphosite on the lumenal PsbQ-2 subunit is not shown in the current view and so is the PsbR phosphoprotein due to its uncertain location. A superimposition of shared (*B*) and unique (*C*) oxidative protein modifications onto RC47 complex. Only modification of core protein subunits that are common to both RC47 and C (D1, D2, and CP47) are shown. Protein subunits are rendered in lighter colors (D1, green; D2, yellow; and CP47, blue) and their corresponding modifications in darker shades. The oxidatively modified amino acid residues are represented by spheres. Some oxidatively modified amino acids contain more than one type of modifications (Tables S1 and S2). The peroxide-induced uniquely modified D1 protein residues found in proximity to the D1-CP43 interface are circled in red. The Mn-cluster, hon-heme iron, bicarbonate ligand of the non-heme iron, P680 special pair, pheophytin, and Q_A_ and Q_B_ quinones are also depicted. The shared and unique oxidative modification of monomeric core are shown in Fig. S4. All oxidative protein modifications depicted are those reported in Fig. 4*B* and listed in Tables S1 and S2. The C2S2M2 and C structures are based on pea PSII structure 5XNM.pdb (12). All oxidatively modified residues shown are fully conserved between *Arabidopsis* and pea.

**Figure 6.**
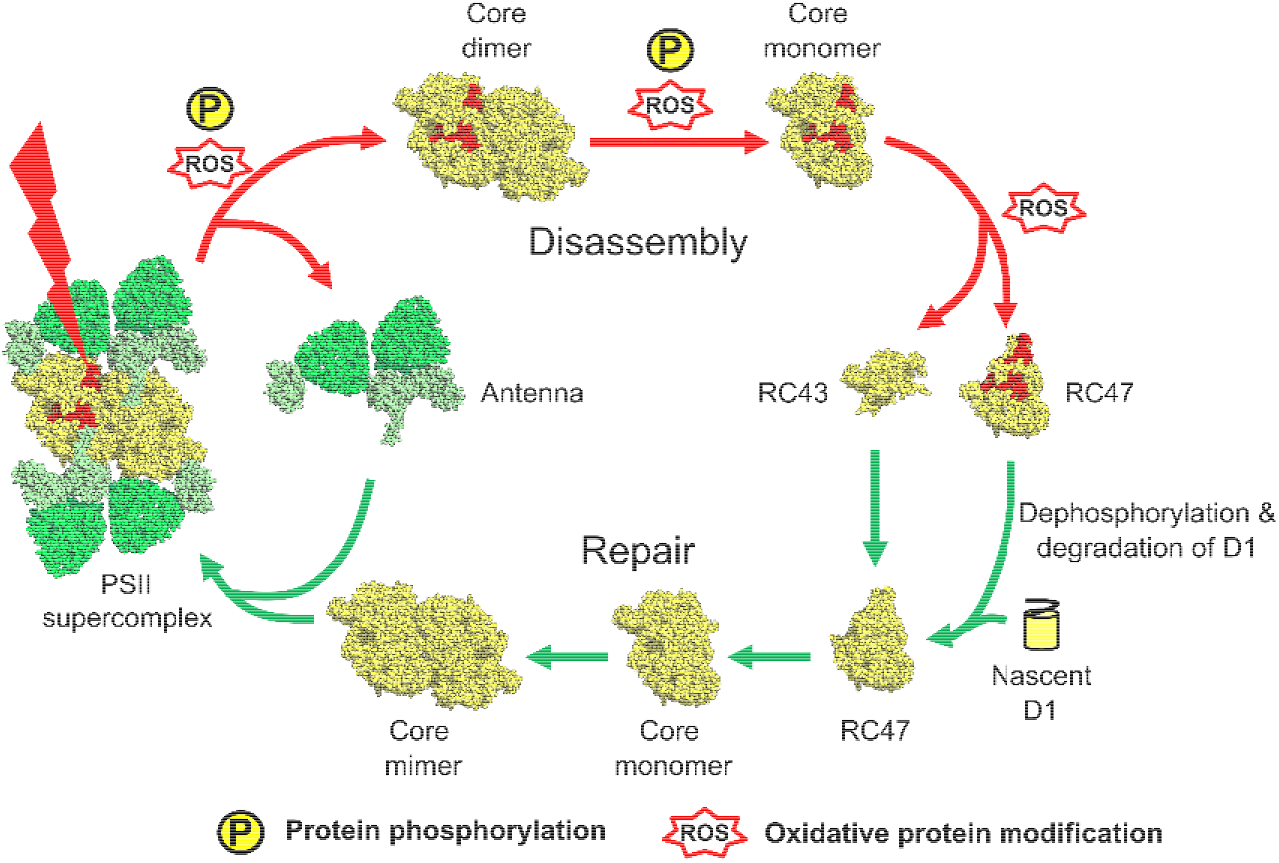
A scheme for plant PSII repair cycle. The damaged D1 protein depicted in red. Phosphorylation and oxidative protein modification are suggested to act synergistically for the sequential and economical disassembly of photodamaged PSII. Phosphorylation may additionally help to prevent the reassembly of unrepaired cores. Oxidative protein modifications may further help the D1 protease to recognize and selectively degrade the damaged D1 proteins.

LHCII and core phosphorylation are readily reversible by unique as well as slightly redundant activities of PSII phosphatases TAP38/PPH1 and PBCP, respectively (22–24). The preferential dephosphorylation of PSII core phosphoproteins by PBCP is likely to be a key event in PSII repair cycle as the efficient degradation of photodamaged D1 by its proteases requires prior D1 dephosphorylation (47). Dephosphorylation may similarly be necessary for the reassembly of repaired reaction center cores into PSII supercomplexes given our (Figs. 1 and 2) and others’ demonstration that core phosphorylation leads to disassembly (25, 26). An implication of the latter is that phosphorylation will prevent reassembly if phosphorylated cores are not swiftly acted up on by PBCP. The glutathione-mediated inhibition of PBCP in high light is consistent with this dual role of phosphorylation in initiating disassembly as well as preventing reassembly of disassembled complexes (48). A phosphorylation-driven pause in reassembly may thus ensure a temporal uncoupling of damage and repair so that only repaired monomeric cores reassemble with the peripheral antenna. This delay in repair and reassembly under photoinhibitory conditions may also prevent continued damage given the smaller absorption cross section of reaction center cores and the quenched nature of detached antenna (49). The altered PSII structural species distribution in kinase and phosphatase mutants and STN8 overexpressor (Fig. 1D) may thus reflect these opposing and kinetically different activities of the kinase and phosphatase. However, the shift in PSII distribution towards more dissembled complexes in *in vitro*-phosphorylated samples (Fig. 2B) strongly implies initiation of PSII disassembly by core phosphorylation as these thylakoids lack PBCP (22, 30). The differing levels of disassembly in light treated ATP plus and minus samples (Fig. 2B) similarly argue for an active role of phosphorylation in triggering PSII disassembly as other phosphorylation-independent disassembly mechanisms are likely to be operating at similar extent in these two thylakoids.

Our analysis of PSII disassembly in kinase and phosphatase mutants and in wild type and *stn7stn8 in vitro* phosphorylated thylakoid samples (Figs. 1 and 2) makes it abundantly clear phosphorylation accounts for only a proportion of disassembly and that other light-dependent mechanisms should be sought for fully explaining PSII disassembly. Mechanisms such as high light-induced LHCII aggregation and changes in thylakoid lipid composition might contribute to PSII disassembly. The high light-induced aggregation of LHCII is driven by the nonpigment-binding PsbS protein (50). It is believed that the protonation of a few lumenal glutamate residues in PsbS induces a change in its conformation, which in turn causes the aggregation of LHCII (51, 52). The aggregated LHCII dissipates absorbed light energy as heat in a photoprotective pathway known as the energy-dependent nonphotochemical quenching (qE). The PsbS-mediated LHCII aggregation could enable the dissociation of the M-trimer from the core as PsbS is thought to be localized in the cleft between the M-trimer and the core (12). Interestingly, a dissociated multimeric antenna complement that comprises M-trimer, CP24, and CP29 has been identified as a site for qE in *Arabidopsis* (49).

While the involvement of processes such as qE in PSII disassembly requires further investigation, here we provide compelling evidence for protein oxidative modification as a phosphorylation-independent mechanism for PSII disassembly (Figs. 3 and 4). Oxidative modification seems especially important for the dissociation of the monomeric cores into submonomeric complexes (Fig. 3*B* and *C*) but it cannot be ruled out whether photooxidative damage acts alone or synergistically with phosphorylation for the disassembly of the peripheral antenna from the core and in core monomerization. The exogenous peroxide-driven disassembly of the *stn7stn8* reaction center core was especially striking (Fig. 3*C*), offering insights into a solely damage-mediated PSII disassembly mechanism in the kinase double mutant. The exact *in vivo* chloroplast concentration of H_2_O_2_ is uncertain, but we demonstrate PSII disassembly even under micromolar peroxide concentrations. Admittedly, some of the higher H_2_O_2_ concentrations used are not physiologically relevant but our aim was to demonstrate a dose-dependency in the disassembly response (Fig. 3*A*). A projection of the shared and unique oxidative protein modifications onto the RC47 complex (Fig. 5*B* and *C*) shows an enrichment of modified amino acids in the vicinity of the Mn-cluster and Q_A_-binding site, as pertinent to the localized nature of the origin and spread of reactive oxygen species from these two cofactors in PSII (1). At the acceptor side of PSII, some of the oxidative modifications are clustered at the interface of D1 and CP43 subunits. These oxidative modifications may weaken the D1-CP43 interaction, allowing the dissociation of the monomeric cores along these subunits. A role for reactive oxygen species in the selective cleavage of the D1 protein has been previously noted (53). This may represent an additional mechanism for the destabilization of the PSII monomeric core. Since phosphoryl groups are removed from D1 prior to its degradation (47), oxidative modifications may instead enable the protease machinery to recognize and selectively degrade the damaged D1. By these mechanisms, oxidative modifications may ensure the disassembly and degradation of only the damaged monomeric cores, sparing undamaged monomeric cores for swift reassembly into PSII supercomplexes. Oxidative protein modification may especially be significant for the disassembly and repair of PSII in organisms such as cyanobacteria and nongreen algae that altogether lack PSII phosphorylation (10). Oxidative protein modification is likely to be the ancient disassembly mechanism of PSII and core protein phosphorylation might have originated later in evolution to impart explicit control over the disassembly and repair processes.

## Materials and Methods

### Plant materials, growth conditions, and isolation of thylakoid membranes

*Arabidopsis thaliana* (Col-0) wild type, *stn8-1, stn7stn8, STN8oe, pbcp-1*, and *pbcp-2* plants were grown from seeds on soil at 21 °C under a white growth light of ∼150 μmol m^-2^ s^-1^ on an 8-hour light and 16-hour dark short-day condition. The knockout mutants were obtained from the Arabidopsis Biological Resource Center (ABRC) confirmed T-DNA insertion mutant collection: SALK_073254 (*stn7-1*), SALK_060869 (*stn8-1*), SALK_127920.31.10.N (*pbcp-1*), and WiscDsLox_359F02 (*pbcp-2*). *STN8oe* was derived from the overexpression of *STN8* coding sequence from a Cauliflower Mosaic Virus (CaMV) 35S promoter in the *stn8-1* mutant background. The moderate light (∼800 μmol m^-2^ s^-1^) illumination employed for kinase and phosphatase mutant characterization was provided by LED light bulbs (Feit Electric). The far-red light used for *in vitro* thylakoid phosphorylation assay of wild type and *stn7stn8* was supplied by Narva 18 W/015 red fluorescent strip lamps wrapped with a layer of plasa red filter (LEE 029), giving ∼J6 μmol m^-2^ s^-1^ at the leaf level. 4-8-week-old plants were used for all experiments.

Thylakoid membranes were isolated in darkness as described previously (54), and resuspended in a small volume of ice-cold storage buffer C (0.1 M Sorbitol, 50 mM HEPES, 10 mM MgCl_2_, 15 mM NaCl, pH 7.5) containing protease inhibitors (1 mg/mL Pefabloc SC, 0.3 μg/mL Leupeptin, 0.2 μg/mL Antipain, and 200 μM PMSF) and 10 mM NaF. Chlorophyll concentration of thylakoid samples was determined in 80% aqueous acetone according to the Porra method (55).

### Chlorophyll fluorescence measurements

For chlorophyll fluorescence induction measurement, fresh thylakoid membranes were isolated from 6-8-week-old *Arabidopsis* plants. Thylakoid suspension was diluted to a final chlorophyll concentration of 10 μg/mL in storage buffer C, containing 150 mM Sorbitol, and mixed with DCMU at a final concentration of 20 μM. The reaction mixture was kept on ice in complete darkness prior to each measurement. Chlorophyll fluorescence was recorded with a JTS-10 spectrophotometer (SpectroLogiX) using a custom script. Briefly, thylakoid samples were excited with a weakly absorbing green LED actinic light (30 μmol m^-2^ s^-1^). The detection pulses were provided by a white LED light passing through a blue filter (470 nm) to preferentially excite chlorophyll a molecules. The measuring photodiode was protected with an RG665 interference filter (Schott, Mainz, Germany) and the reference diode with a BG39 filter (Schott, Mainz, Germany).

### Polyacrylamide gel electrophoresis

For immunoblot analysis, thylakoid protein samples were separated by an 11.5% SDS-Urea PAGE gel system. BN-PAGE analysis of thylakoid membranes was performed essentially as described earlier (56). Briefly, freshly isolated thylakoid samples equivalent to 50 μg of chlorophyll were washed twice in TMK buffer (10 mM Tris-HCl, pH 6.8, 10 mM MgCl_2_, 20 mM KCl) at 5,000 rpm at 4 °C on a benchtop centrifuge at a total volume of 300 μL. Supernatant was discarded and washed thylakoids were resuspended in 60 μL of ACA buffer (750 mM ε-Aminocaproic Acid, 50 mM Bis-Tris, pH 7, 5 mM EDTA, 50 mM NaCl) and mixed with 6 μL of freshly prepared 10% α-DDM to a final concentration of 0.9% α-DDM, briefly vortexed, and incubated on ice for 10 minutes. Samples were centrifuged at 13,000 rpm for 10 minutes at 4 °C on a benchtop centrifuge, and supernatant was mixed with 6 μL loading buffer (750 mM ε-aminocaproic acid and 5% (w/v) pure Coomassie Brilliant Blue G (B0770, Sigma)) and loaded onto a 6-12% gradient polyacrylamide gel. Gels were run at 4 °C for 16 hours overnight with blue cathode buffer (50 mM Tricine, 15 mM Bis-Tris pH 7.0, supplemented w/ 0.02% Coomassie G-250) at 60 V. After the overnight run, blue cathode buffer was exchanged with colorless cathode buffer lacking Coomassie G-250, and the gel was run at 250 V for an additional 2-4 hours at 4 °C. For analysis of oxidative modification, BN-PAGE gels were polymerized with 10 μM flavin mononucleotide in the presence of long-wavelength UV light to avoid artifactual oxidative protein modification from ammonium persulfate, and the gel was run with cathode buffer containing 100 mM sodium thioglycolate.

### Immunoblotting

Each thylakoid sample was diluted with breaking buffer (130 mM Tris-HCl, pH 6.8, 16% (w/v) glycerol, 4.6% (w/v) SDS) and mixed with 5x Laemmli buffer, and a sample volume equivalent to 0.5 μg chlorophyll was loaded onto an SDS-PAGE gel. The gel was run at 20 mA constant current for 3 hours, and the proteins transferred onto a PVDF membrane at 20 V for 1 hour. Upon transfer, the membrane was washed three times for 5 minutes each in TBS-Tween (0.1%), blocked in blocking buffer overnight, and incubated with primary antibody for 1 hour and HRP-conjugated secondary antibody for 1 hour. Immunoreactive bands were visualized on a ChemiDoc MP imager (Bio-Rad) using a chemiluminescence detection reagent (ECL Clarity, Bio-Rad). Primary antibodies used are anti-phosphothreonine polyclonal antibody (9381, Cell Signaling) and anti-D1 polyclonal antibody (AS05084, Agrisera). The secondary antibody is an anti-rabbit IgG (AS09602, Agrisera).

For immunoblot analysis of BN-PAGE gels, gels were first incubated in a denaturing solution consisting of 1x Laemmli buffer, devoid of bromophenol blue, for 30 minutes at room temperature, followed by Towbin buffer for 1 hour at room temperature. The proteins from the gel were transferred to a PVDF membrane at 180 mA for 2 hours, and washed three times for 5 minutes each in a destain solution (10% Acetic Acid, 45% MeOH) prior to blocking.

### In vitro phosphorylation of thylakoids

6-week-old wild type and *stn7stn8* plants were moved to darkness for 16 hours, followed by 2 hours of far-red light illumination. Thylakoids were promptly isolated from far-red-treated plants in darkness, and chlorophyll concentration was determined. Thylakoid membranes were diluted to 0.5 μg/μL chlorophyll concentration in phosphorylation buffer (20 mM Tris-HCl pH 7.5, 5 mM MgCl_2_, 1 mM MnCl_2_ and 20 mM NaF) and mixed with 25 μM ATP or buffer. In vitro phosphorylation was initiated by illumination of the samples with a white LED light (∼150 μmol m^-2^ s^-1^) for 1 hour at 21 °C. Aliquots of far-red-treated diluted thylakoid samples, without ATP, were kept in darkness for 1 hour at 21 °C and used as control. For western blot analysis, thylakoid samples equivalent to 0.5 μg chlorophyll were immediately mixed with 5x Laemmli buffer and loaded onto an 11.5% SDS-Urea PAGE gel. For blue-native PAGE analysis, thylakoid samples equivalent to 50 μg chlorophyll were withdrawn and processed for BN-PAGE as described earlier.

### Exogenous ROS treatment

Thylakoid membranes equivalent to 50 μg chlorophyll were diluted to 0.3 μg chlorophyll/μL in storage buffer C, containing protease inhibitors and 10 mM NaF, and mixed with varying concentrations of H_2_O_2_ at 10% of the total reaction volume. Isolated PSII core complexes were treated essentially in the same manner but starting with a total chlorophyll content of 10 μg. Reactions were allowed to proceed in darkness at room temperature for 30 minutes or up to 1 hour, depending on the experiment.

### Isolation of PSII core complexes

PSII core particles were isolated as reported in van Leeuwen et al. (57), with a few minor modifications. To prepare BBY membranes (named after Berthold, Babcock and Yocum (58)), *Arabidopsis* rosettes were washed and homogenized in B1 solution (20 mM Tricine, pH 7.8, 0.4 M NaCl, 2 mM MgCl_2_, 1 mM 6-aminocaproic acid, 0.2 mM Benzamidine) and passed through 8 layers of cheesecloth and 1 layer of Miracloth. Pellets were collected by centrifugation at 1400 x g at 4 °C for 10 minutes, and resuspended in B2 solution (20 mM Tricine, pH 7.8, 0.15 M NaCl, 5 mM MgCl_2_, 1 mM 6-aminocaproic acid, 0.2 mM Benzamidine). The resulting membrane suspension was centrifuged at 4000 x g at 4 °C for 10 minutes, and the pellet was resuspended in B3 solution (20 mM HEPES, pH 7.5, 15 mM NaCl, 5 mM MgCl_2_). The membranes were subsequently pelleted by centrifugation at 6000 x g and resuspended again in B3 solution to a 2 mg/mL chlorophyll concentration, and Triton X-100 was added dropwise to a final concentration of 25 mg/mg chlorophyll with slow constant stirring for 15 minutes. Triton-solubilized thylakoids were centrifuged at 40000 x g for 25 minutes at 4 °C, pellet resuspended in B3 solution, and spun at 2000 x g for 5 minutes at 4 °C to pellet out starch granules. Membranes from the remaining supernatant were pelleted once again by centrifugation at 40000 x g for 25 minutes at 4 °C and stored in BBY storage buffer (20 mM HEPES, pH 7.5, 0.4 M Sorbitol, 15 mM NaCl, 5 mM MgCl_2_) containing protease inhibitors and 10 mM NaF at -80 °C for further use.

BBY membranes were further washed in Solution B4 (20 mM MES, pH 6, 15 mM NaCl, 5 mM CaCl_2_, 0.4 M Sucrose), pelleted by centrifugation at 40000 x g for 25 minutes at 4 °C, and subsequently resuspended in BTS400 (20 mM Bis-Tris, pH 6.5, 20 mM MgCl_2_, 5 mM CaCl_2_, 10 mM MgSO_4_, 0.4 mM Sucrose, 0.03% β-DM). The membranes were adjusted to a 2 mg/mL chlorophyll concentration and solubilized by addition of 1/7^th^ volume of 10% w/v β-DM to a final detergent concentration of 1.25% w/v. The membrane-detergent suspension was mixed by end-over-end inversion for 10 minutes, and insoluble material was pelleted at 40000 x g for 20 minutes at 4 °C. Solubilized membranes were applied to a Q-Sepharose FF (Pharmacia Biotech) column pre-equilibrated with BTS400. The column was washed with BTS400 until the eluate became clear, and PSII core complexes were eluted with several column volumes of BTS400 containing 75 mM MgSO_4_. Purified PSII core complexes were further concentrated by a 30 kDa MWCO filter (Amicon) and desalted into storage buffer C containing protease inhibitors and 10 mM NaF.

### LC-MS/MS

For oxidative modification analysis, bands corresponding to C and RC47 were excised from the blue native gels using a new razor, reduced in 10 mM DTT at 55 °C for 1 hour, and alkylated in 55 mM iodoacetamide in darkness for 45 minutes. In-gel digestion of proteins was performed at 37 °C overnight using 2.5 μg Trypsin/Lys-C (Promega). Samples were further digested with 2.5 μg Chymotrypsin (Promega) at 25 °C for 4 hours. For phosphopeptide enrichment, proteins were extracted from thylakoid samples by heating at 80 °C for 10 minutes in 40 mM Tris-HCl (pH 8), 40 mM DTT, and 4% SDS, followed by chloroform-methanol precipitation and resuspension in 8 M urea, 20 mM Tris-HCl (pH 8), and 3 mM EDTA. Trypsin digestion was performed essentially as above with the omission of chymotrypsin. Phosphopeptides were enriched using PolyMac beads according to the manufacturer’s instructions. Peptides were desalted with a C18 spin column according to manufacturer’s instructions (The Nest Group). Peptides were dried and resuspended in 3% acetonitrile/0.1% formic acid and peptide concentration was measured by BCA assay (Pierce) and equal amounts of peptides were run on LC-MS/MS as described previously (54).

For oxidative modifications analysis, raw files were searched using MaxQuant (1.6.14) against a custom FASTA library containing D1, D2, and CP47 proteins. Unless explicitly stated, all parameters were left in their default settings. Digestion enzymes included Trypsin/P, Chymotrypsin, and Lys-C, with a max number of missed cleavages set to 2. Oxidation (M) and Acetylation (N-term) were set as variable modifications, and Carbamidomethylation (C) was set as a fixed modification, with max number of modifications per peptide set to 5. Searches for oxidative modifications were performed by addition of the respective variable modifications with specificities and monoisotopic masses derived from Unimod. Main search peptide tolerance was set to 10 ppm, minimum peptide length was set to 7, and PSM FDR was set to 1%. Modified peptides were analyzed from the modificationSites text file, and filtered for contaminants. Only high confidence modifications were considered with an amino acid localization probability of ≥ 0.75 and a mass error of ≤ 5 ppm. Additionally, only modifications detected in all three biological replicates were considered for this analysis.

For phosphoproteome analysis, phosphopeptides were searched in MaxQuant (1.6.14) against the TAIR10 protein FASTA database using Trypsin/P, and Lys-C as digestion enzymes with a max number of missed cleavages set to 2. Oxidation (M), Acetylation (N-term), and Phosphorylation (ST) were set as variable modifications, and Carbamidomethylation (C) was set as a fixed modification, with max number of modifications per peptide set to 5. Main search peptide tolerance was set to 10 ppm, and PSM FDR was set to 1%.

## Acknowledgments

This research is supported by grants to S.P. from the United States Department of Energy (DOE) (DE-SC0020639) and the United States Department of Agriculture-National Institute of Food and Agriculture (USDA-NIFA) (Hatch: 1013608). S.D.M. acknowledges USDA-NIFA for a EWD predoctoral fellowship (2021-67034-35183). We thank Dr. Uma Aryal (Purdue Proteomics Core) for help with running of mass spectrometry samples and Dr. Iskander M. Ibrahim for discussions.

## Notes

### Competing Interest Statement

The authors have declared no competing interest.

## References

1. R. Kale, et al., Amino acid oxidation of the D1 and D2 proteins by oxygen radicals during photoinhibition of Photosystem II. Proc. Natl. Acad. Sci. U. S. A. 114, 2988–2993 (2017).

2. G. A. Davis, et al., Limitations to photosynthesis by proton motive force-induced photosystem II photodamage. Elife 5 10.7554/eLife.16921 (2016).

3. A. K. Mattoo, U. Pick, H. Hoffman-Falk, M. Edelman, The rapidly metabolized 32,000-dalton polypeptide of the chloroplast is the “proteinaceous shield” regulating photosystem II electron transport and mediating diuron herbicide sensitivity. Proc. Natl. Acad. Sci. U. S. A. 78, 1572–1576 (1981).

4. A. K. Mattoo, H. Hoffman-Falk, J. B. Marder, M. Edelman, Regulation of protein metabolism: Coupling of photosynthetic electron transport to in vivo degradation of the rapidly metabolized 32-kilodalton protein of the chloroplast membranes. Proc. Natl. Acad. Sci. U. S. A. 81, 1380–1384 (1984).

5. D. J. Kyle, I. Ohad, C. J. Arntzen, Membrane protein damage and repair: Selective loss of a quinone-protein function in chloroplast membranes. Proc. Natl. Acad. Sci. U. S. A. 81, 4070–4074 (1984).

6. E. M. Aro, I. Virgin, B. Andersson, Photoinhibition of Photosystem II. Inactivation, protein damage and turnover. Biochim. Biophys. Acta 1143, 113–134 (1993).

7. P. J. Nixon, F. Michoux, J. Yu, M. Boehm, J. Komenda, Recent advances in understanding the assembly and repair of photosystem II. Ann. Bot. 106, 1–16 (2010).

8. S. Järvi, M. Suorsa, E.-M. Aro, Photosystem II repair in plant chloroplasts — Regulation, assisting proteins and shared components with photosystem II biogenesis. Biochim. Biophys. Acta - Bioenerg. 1847, 900–909 (2015).

9. A. Melis, Photosystem-II damage and repair cycle in chloroplasts: what modulates the rate of photodamageJ? Trends Plant Sci. 4, 130–135 (1999).

10. V. M. Johnson, H. B. Pakrasi, Advances in the understanding of the lifecycle of photosystem II. Microorg. 10, 836 (2022).

11. H. Kirchhoff, Structural constraints for protein repair in plant photosynthetic membranes. Plant Signal. Behav. 8, e23634 (2013).

12. X. Su, et al., Structure and assembly mechanism of plant C2S2M2-type PSII-LHCII supercomplex. Science 357, 815–820 (2017).

13. P. Cao, et al., Structure, assembly and energy transfer of plant photosystem II supercomplex. Biochim. Biophys. Acta - Bioenerg. 1859, 633–644 (2018).

14. X. Wei, et al., Structure of spinach photosystem II–LHCII supercomplex at 3.2 Åresolution. Nature 534, 69–74 (2016).

15. T. K. Goral, et al., Visualizing the mobility and distribution of chlorophyll proteins in higher plant thylakoid membranes: effects of photoinhibition and protein phosphorylation. Plant J. 62, 948–959 (2010).

16. M. Herbstova, S. Tietz, C. Kinzel, M. V Turkina, H. Kirchhoff, Architectural switch in plant photosynthetic membranes induced by light stress. Proc. Natl. Acad. Sci. U. S. A. 109, 20130–20135 (2012).

17. N. Depege, S. Bellafiore J. D. Rochaix, Rote of chloroplast protein kinase Stt7 in LHCII phosphorylation and state transition in Chlamydomonas. Science 299, 1572–1575 (2003).

18. S. Bellafiore, F. Barneche, G. Peltier, J. D. Rochaix, State transitions and light adaptation require chloroplast thylakoid protein kinase STN7. Nature 433, 892–895 (2005).

19. V. Bonardi, et al., Photosystem II core phosphorylation and photosynthetic acclimation require two different protein kinases. Nature 437, 1179–1182 (2005).

20. J. D. Rochaix, Role of thylakoid protein kinases in photosynthetic acclimation. FEBS Lett. 581, 2768–2775 (2007).

21. P. Pesaresi, M. Pribil, T. Wunder, D. Leister, Dynamics of reversible protein phosphorylation in thylakoids of flowering plants: the roles of STN7, STN8 and TAP38. Biochim Biophys Acta 1807, 887–896 (2010).

22. I. Samol, et al., Identification of a photosystem II phosphatase involved in light acclimation in Arabidopsis. Plant Cell 24, 2596–2609 (2012).

23. M. Pribil, P. Pesaresi, A. Hertle, R. Barbato, D. Leister, Role of plastid protein phosphatase TAP38 in LHCII dephosphorylation and thylakoid electron flow. PLoS Biol 8, e1000288 (2010).

24. A. Shapiguzov, et al., The PPH1 phosphatase is specifically involved in LHCII dephosphorylation and state transitions in Arabidopsis. Proc. Natl. Acad. Sci. U. S. A. 107, 4782–4787 (2010).

25. M. Tikkanen, M. Nurmi, S. Kangasjarvi, E. M. Aro, Core protein phosphorylation facilitates the repair of photodamaged photosystem II at high light. Biochim. Biophys. Acta 1777, 1432–1437 (2008).

26. R. Fristedt, A. V Vener, High light induced disassembly of photosystem II supercomplexes in Arabidopsis requires STN7-dependent phosphorylation of CP29. PLoS One 6, e24565 (2011).

27. S. Puthiyaveetil, H. Kirchhoff, A phosphorylation map of the photosystem II supercomplex C2S2M2. Front. Plant Sci. 4, 459 (2013).

28. P. Longoni, I. Samol M. Goldschmidt-Clermont, The kinase STATE TRANSITION 8 phosphorylates light harvesting complex II and contributes to light acclimation in Arabidopsis thaliana. Front. Plant Sci. 10, 1156 (2019).

29. T. Wunder, et al., The major thylakoid protein kinases STN7 and STN8 revisited: effects of altered STN8 levels and regulatory specificities of the STN kinases. Front. Plant Sci. 4, 417 (2013).

30. S. Puthiyaveetil, et al., Significance of the Photosystem II Core Phosphatase PBCP for Plant Viability and Protein Repair in Thylakoid Membranes. Plant Cell Physiol. 55, 1245– 1254 (2014).

31. F. Rappaport, et al., On the advantages of using green light to study fluorescence yield changes in leaves. Biochim. Biophys. Acta - Bioenerg. 1767, 56–65 (2007).

32. M. Pietrzykowska, et al., The light-harvesting chlorophyll a/b binding proteins Lhcb1 and Lhcb2 play complementary roles during state transitions in Arabidopsis. Plant Cell 26, 3646–3660 (2014).

33. M. Suorsa, et al., Light acclimation involves dynamic re-organization of the pigment-protein megacomplexes in non-appressed thylakoid domains. Plant J. 84, 360–373 (2015).

34. R. Kouřil, et al., High-light vs. low-light: effect of light acclimation on photosystem II composition and organization in Arabidopsis thaliana. Biochim. Biophys. Acta 1827, 411– 419 (2013).

35. V. Krynická, S. Shao, P. J. Nixon, J. Komenda, Accessibility controls selective degradation of photosystem II subunits by FtsH protease. Nat. Plants 1, 1–6 (2015).

36. J. Nickelsen, B. Rengstl, Photosystem II assembly: from cyanobacteria to plants. Annu. Rev. Plant Biol. 64, 609–635 (2013).

37. I. Baroli, A. Melis, Photoinhibitory damage is modulated by the rate of photosynthesis and by the photosystem II light-harvesting chlorophyll antenna size. Planta 205, 288–296 (1998).

38. M. Tikkanen, E. M. Aro, Thylakoid protein phosphorylation in dynamic regulation of photosystem II in higher plants. Biochim. Biophys. Acta 1817, 232–238 (2012).

39. E. Rintamaki, P. Martinsuo, S. Pursiheimo, E. M. Aro, Cooperative regulation of light-harvesting complex II phosphorylation via the plastoquinol and ferredoxin-thioredoxin system in chloroplasts. Proc. Natl. Acad. Sci. U. S. A. 97, 11644–11649 (2000).

40. J. F. Allen, J. Bennett, K. E. Steinback, C. J. Arntzen, Chloroplast Protein Phosphorylation Couples Plastoquinone Redox State to Distribution of Excitation-Energy between Photosystems. Nature 291, 25–29 (1981).

41. J. Bennett, Phosphorylation of Chloroplast Membrane Polypeptides. Nature 269, 344–346 (1977).

42. M. Suorsa, et al., Protein assembly of photosystem II and accumulation of subcomplexes in the absence of low molecular mass subunits PsbL and PsbJ. Eur. J. Biochem. 271, 96– 107 (2004).

43. X. Yi, S. R. Hargett, L. K. Frankel, T. M. Bricker, The PsbQ protein is required in Arabidopsis for photosystem II assembly/stability and photoautotrophy under low light conditions. J. Biol. Chem. 281, 26260–26267 (2006).

44. K. Ido, et al., Cross-linking evidence for multiple interactions of the PsbP and PsbQ proteins in a higher plant photosystem II supercomplex. J. Biol. Chem. 289, 20150–20157 (2014).

45. U. Ljungberg, H. E. Åkerlung, C. Larsson, B. Andersson, Identification of polypeptides associated with the 23 and 33 kDa proteins of photosynthetic oxygen evolution. Biochim. Biophys. Acta - Bioenerg. 767, 145–152 (1984).

46. M. Suorsa, et al., PsbR, a missing link in the assembly of the oxygen-evolving complex of plant photosystem II. J. Biol. Chem. 281, 145–150 (2006).

47. E. Rintamaki, et al., Differential D1 dephosphorylation in functional and photodamaged photosystem II centers: dephosphorylation is a prerequisite for degradation of damaged D1. J. Biol. Chem. 271, 14870–14875 (1996).

48. X. Liu, J. Chai, X. Ou, M. Li, Z. Liu, Structural insights into substrate selectivity, catalytic mechanism, and redox regulation of rice photosystem II core phosphatase. Mol. Plant 12, 86–98 (2018).

49. N. Betterle, et al., Light-induced dissociation of an antenna hetero-oligomer is needed for non-photochemical quenching induction. J. Biol. Chem. 284, 15255–15266 (2009).

50. X. P. Li, et al., A pigment-binding protein essential for regulation of photosynthetic light harvesting. Nature 403, 391–395 (2000).

51. X.-P. Li, et al., Regulation of photosynthetic light harvesting involves intrathylakoid lumen pH sensing by the PsbS protein. J. Biol. Chem. 279, 22866–22874 (2004).

52. A. V Ruban, Evolution under the sun: optimizing light harvesting in photosynthesis. J. Exp. Bot. 66, 7–23 (2015).

53. M. Miyao, M. Ikeuchi, N. Yamamoto, T. Ono, Specific degradation of the D1 protein of photosystem II by treatment with hydrogen peroxide in darkness: implications for the mechanism of degradation of the D1 protein under illumination. Biochemistry 34, 10019– 10026 (1995).

54. S. D. McKenzie, I. M. Ibrahim, U. K. Aryal, S. Puthiyaveetil, Stoichiometry of protein complexes in plant photosynthetic membranes. Biochim. Biophys. Acta - Bioenerg. 1861, 148141 (2020).

55. R. J. Porra, The chequered history of the development and use of simultaneous equations for the accurate determination of chlorophylls a and b. Photosynth. Res. 73, 149–156 (2002).

56. H. Koochak, S. Puthiyaveetil, D. L. Mullendore, M. Li, H. Kirchhoff, The structural and functional domains of plant thylakoid membranes. Plant J. 97, 412–429 (2019).

57. P. J. van Leeuwen, M. C. Nieveen, E. J. van de Meent, J. P. Dekker, H. J. van Gorkom, Rapid and simple isolation of pure photosystem II core and reaction center particles from spinach. Photosynth. Res. 28, 149–153 (1991).

58. D. A. Berthold, G. T. Babcock, C. F. Yocum A highly resolved, oxygen-evolving photosystem II preparation from spinach thylakoid membranes. FEBS Lett. 134, 231–234 (1981).

59. R. Croce, H. van Amerongen, The complex that conquered the land. Science 357, 752 (2017).

