## Supporting Information for "Protein phosphorylation and oxidative protein modification mediate plant photosystem II disassembly and repair"

**Supporting figures**

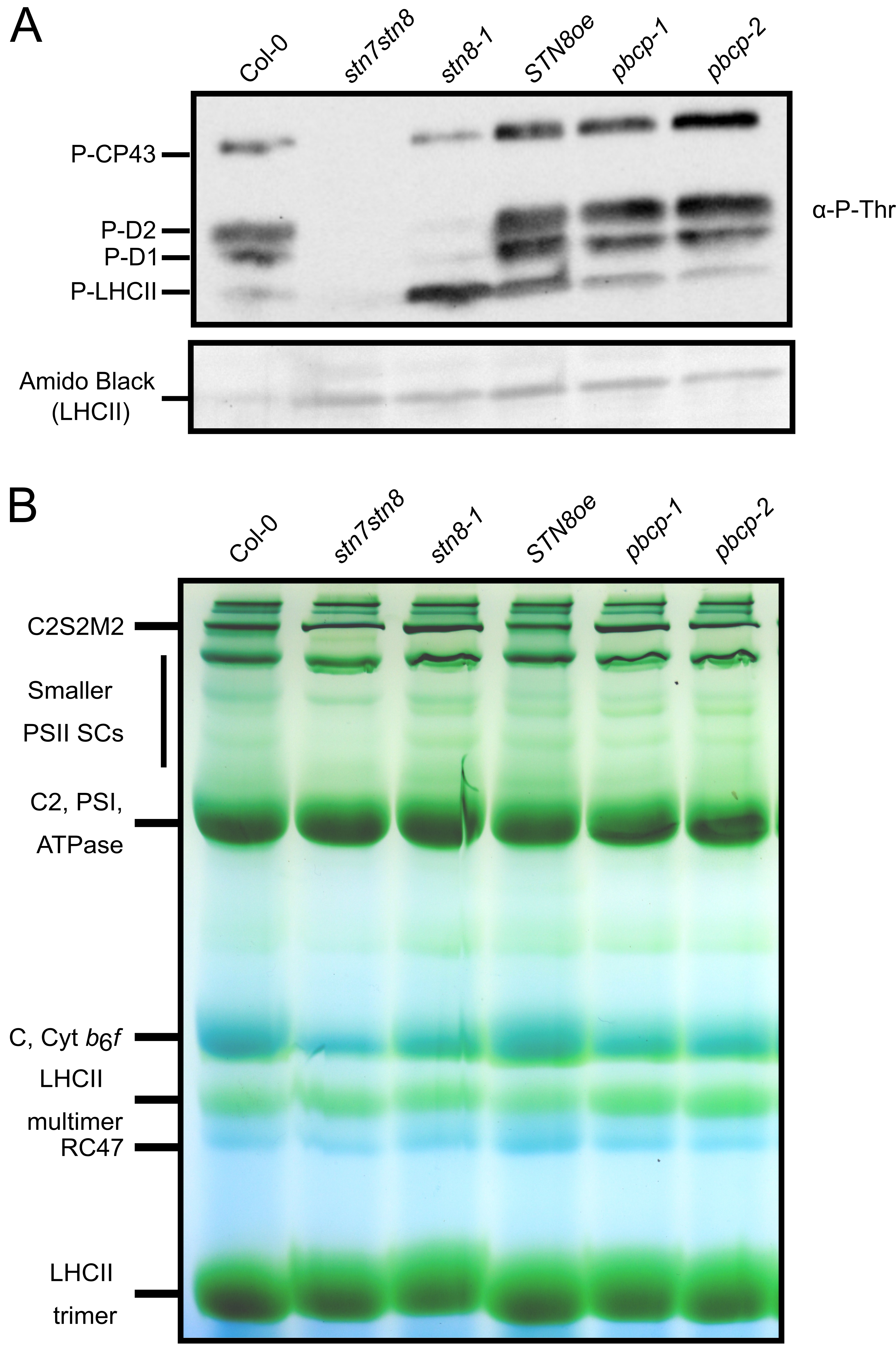

**Figure S1.** PSII assembly status as a function of core phosphorylation level in moderate light. (*A*) A phosphothreonine immunoblot of moderate light-treated thylakoid samples from wild type (Col-0) and mutant plants. Positions of four major PSII phosphoproteins are indicated on the left. The amido black staining of the corresponding PVDF membrane is given as a loading control. (B) A representative blue native gel of moderate light-treated wild type and mutant thylakoid samples. Positions of major thylakoid protein complexes are indicated on the left.

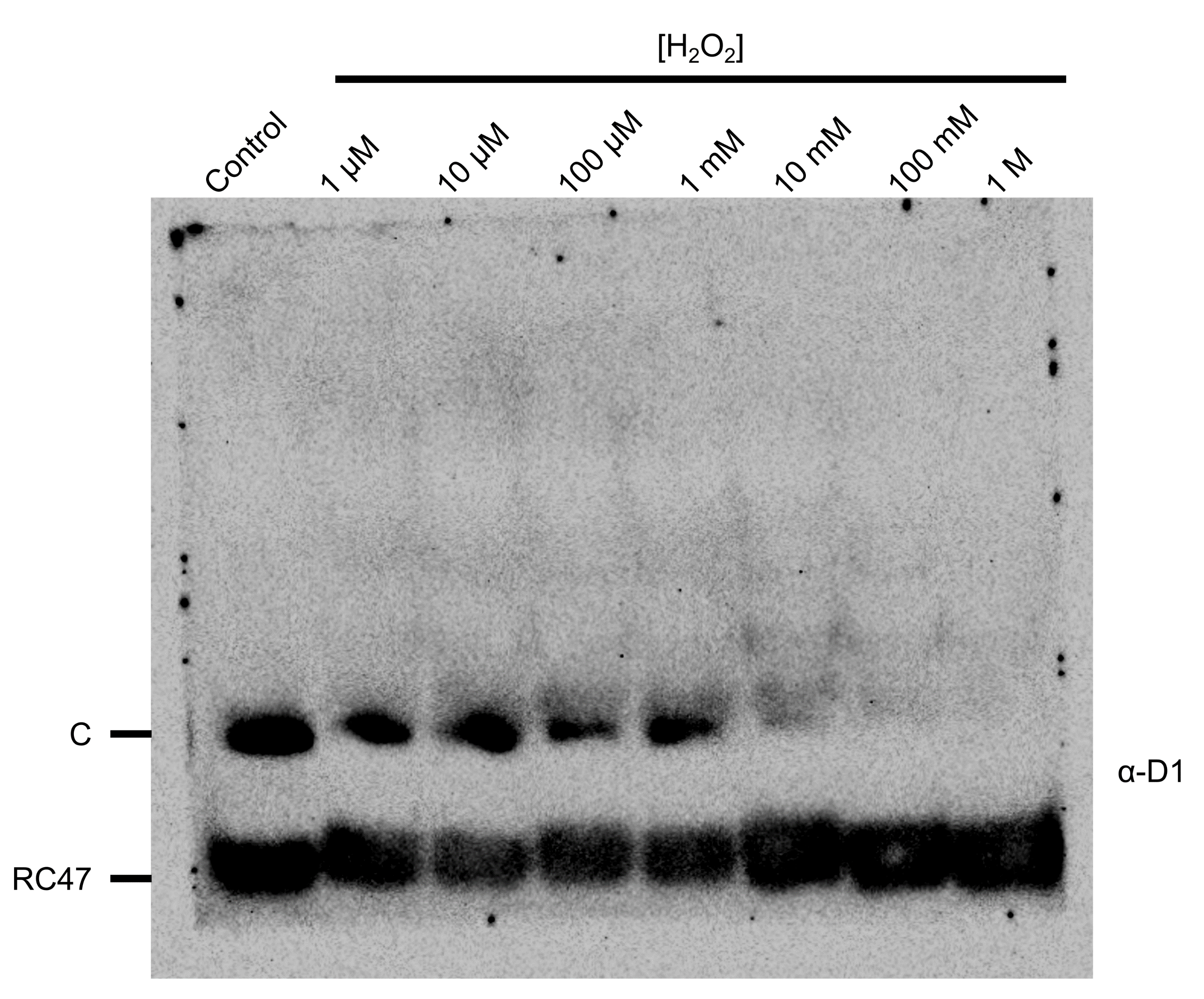

**Figure S2.** H_2_O_2_ treatment results in preferential disassembly of monomeric cores into RC47. An uncropped D1 immunoblot of a blue native gel containing peroxide-treated wild type thylakoid samples. The D1 antibody appears to recognize only C and RC47 complexes but not C2 or PSII supercomplexes. The cropped version of this figure is presented as Fig. 3*B*.

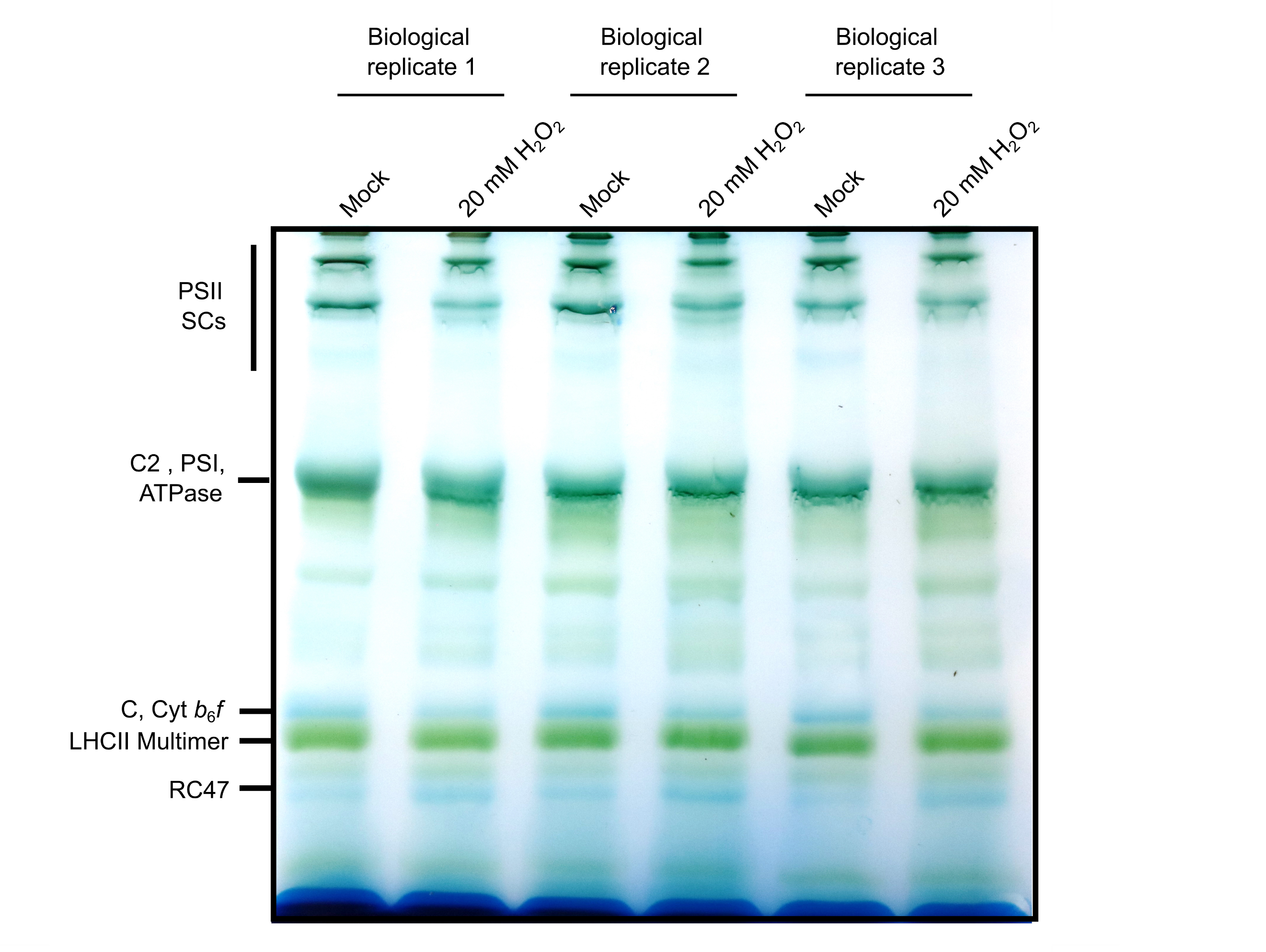

**Figure S3.** A non-oxidizing blue native gel of peroxide-treated wild type thylakoids. Major PSII structural species are indicated on the left. Effects of peroxide treatment on the abundance of C and RC47 complexes in three independent biological replicates are shown. The cropped version of this figure with just one biological replicate is presented as Fig. 4*A*.

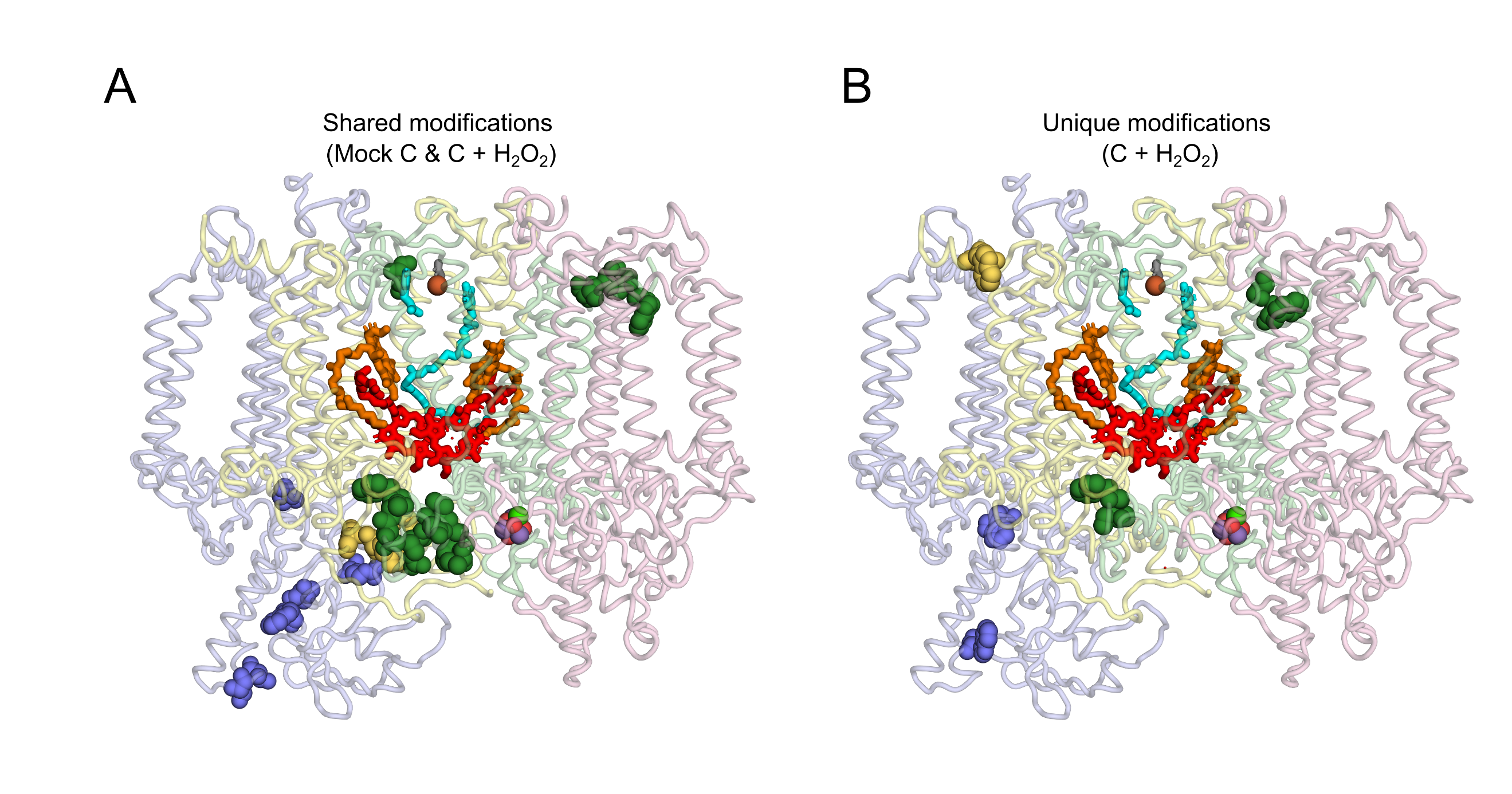

**Figure S4.** A projection of shared (*A*) and unique (*B*) oxidative modifications onto monomeric core complex. Oxidatively modified residues are represented by spheres. Only modifications of D1, D2, and CP47 are shown since these three core subunits are common to both C and RC47 complexes. CP43 subunit is depicted in pink but no modifications are shown. Subunits are colored in lighter shades (D1, green; D2, yellow; and CP47, blue;) and their modifications in darker shades. The Mn-cluster, non-heme iron, bicarbonate ligand of non-heme iron, P680 special pair of chlorophylls, pheophytin, and the Q_A_ and Q_B_ quinones are also shown.

**Table S1.** Shared and unique oxidative modification of RC47 complex under H_2_O_2_ and mock

| **Gene (Protein)** | **Modification probabilities** | **Amino acid** | **Position** | **Modification** | **Reproducible**  **RC47 + H_2_O_2_** | **Reproducible**  **RC47 + Mock** |
| --- | --- | --- | --- | --- | --- | --- |
| ATCG00020.1 (PSBA/D1) | ESES(1)LWGR(1) | S | 12 | Mono-Oxidation | TRUE^1^ | TRUE |
| ATCG00020.1 (PSBA/D1) | ESESLW(0.991)GR(0.009) | W | 14 | Di-Oxidation | TRUE | TRUE |
| ATCG00020.1 (PSBA/D1) | ESES(1)LWGR(1) | R | 16 | Mono-Oxidation | TRUE | TRUE |
| ATCG00020.1 (PSBA/D1) | FCN(1)WITSTENR | N | 19 | Mono-Oxidation | TRUE | TRUE |
| ATCG00020.1 (PSBA/D1) | FCNW(1)ITSTENR | W | 20 | Oxolactone | FALSE^2^ | TRUE |
| ATCG00020.1 (PSBA/D1) | FCNW(1)ITSTENR | W | 20 | Di-Oxidation | TRUE | TRUE |
| ATCG00020.1 (PSBA/D1) | FCNWI(0.996)TSTE(0.004)NR | I | 21 | Carbonylation | FALSE | TRUE |
| ATCG00020.1 (PSBA/D1) | FCN(0.003)WIT(0.925)S(0.923)T(0.148)EN(0.001)R | T | 22 | Mono-Oxidation | TRUE | TRUE |
| ATCG00020.1 (PSBA/D1) | FCN(0.003)WIT(0.925)S(0.923)T(0.148)EN(0.001)R | S | 23 | Mono-Oxidation | TRUE | TRUE |
| ATCG00020.1 (PSBA/D1) | EW(1)ELSFR | W | 131 | Kynurenine | TRUE | TRUE |
| ATCG00020.1 (PSBA/D1) | EW(1)ELSFR | W | 131 | Di-Oxidation | TRUE | FALSE |
| ATCG00020.1 (PSBA/D1) | EWELS(0.999)FR(0.001) | S | 134 | Mono-Oxidation | TRUE | TRUE |
| ATCG00020.1 (PSBA/D1) | LGM(0.986)R(0.986)P(0.986)WIAV(0.042)AY | M | 139 | Mono-Oxidation | TRUE | FALSE |
| ATCG00020.1 (PSBA/D1) | LGM(0.986)R(0.986)P(0.986)WIAV(0.042)AY | R | 140 | Mono-Oxidation | TRUE | FALSE |
| ATCG00020.1 (PSBA/D1) | LGM(0.986)R(0.986)P(0.986)WIAV(0.042)AY | P | 141 | Mono-Oxidation | TRUE | FALSE |
| ATCG00020.1 (PSBA/D1) | LGMRPW(1)IAVAY | W | 142 | Tri-Oxidation | FALSE | TRUE |
| ATCG00020.1 (PSBA/D1) | ETTENESAN(0.003)EGYR(0.997) | R | 238 | Mono-Oxidation | TRUE | TRUE |
| ATCG00020.1 (PSBA/D1) | N(0.986)IV(0.014)AAHGYFGR | N | 247 | Mono-Oxidation | TRUE | FALSE |
| ATCG00020.1 (PSBA/D1) | VIN(1)T(1)WADIINR | N | 315 | Mono-Oxidation | TRUE | TRUE |
| ATCG00020.1 (PSBA/D1) | VIN(1)T(1)WADIINR | T | 316 | Mono-Oxidation | TRUE | TRUE |
| ATCG00020.1 (PSBA/D1) | VINTW(1)ADIINR | W | 317 | Kynurenine | TRUE | TRUE |
| ATCG00020.1 (PSBA/D1) | VINTW(1)ADIINR | W | 317 | Oxolactone | TRUE | TRUE |
| ATCG00020.1 (PSBA/D1) | VINTW(1)ADIINR | W | 317 | Hydroxykynurenine | TRUE | FALSE |
| ATCG00020.1 (PSBA/D1) | VINTW(1)ADIINR | W | 317 | Di-Oxidation | TRUE | TRUE |
| ATCG00020.1 (PSBA/D1) | VINTW(1)ADIINR | W | 317 | Tri-Oxidation | TRUE | TRUE |
| ATCG00020.1 (PSBA/D1) | V(0.012)I(0.012)NTWADI(0.877)I(0.098)NR(0.002) | I | 320 | Carbonylation | TRUE | FALSE |
| ATCG00020.1 (PSBA/D1) | VIN(0.003)T(0.003)WADIIN(0.997)R(0.997) | N | 322 | Mono-Oxidation | TRUE | TRUE |
| ATCG00020.1 (PSBA/D1) | VIN(0.003)T(0.003)WADIIN(0.997)R(0.997) | R | 323 | Mono-Oxidation | TRUE | TRUE |
| ATCG00020.1 (PSBA/D1) | ANL(0.996)GME(0.003)V(0.001)MHER | L | 326 | Carbonylation | TRUE | TRUE |
| ATCG00020.1 (PSBA/D1) | ANLGM(0.965)EVM(0.035)HER | M | 328 | Aspartic semialdehyde | FALSE | TRUE |
| ATCG00020.1 (PSBA/D1) | ANLGM(1)EVMHER | M | 328 | Mono-Oxidation | TRUE | TRUE |
| ATCG00020.1 (PSBA/D1) | AN(0.001)LGMEV(0.995)MHER(0.004) | V | 330 | Mono-Oxidation | TRUE | TRUE |
| ATCG00020.1 (PSBA/D1) | ANLGM(0.18)EVM(0.82)HER | M | 331 | Aspartic semialdehyde | FALSE | TRUE |
| ATCG00020.1 (PSBA/D1) | ANLGM(0.98)EV(0.026)M(0.993)HER(0.001) | M | 331 | Mono-Oxidation | TRUE | TRUE |
| ATCG00270.1 (PSBD/D2) | DLFDIM(1)DDWLR | M | 19 | Mono-Oxidation | TRUE | FALSE |
| ATCG00270.1 (PSBD/D2) | DIMDDW(0.998)LR(0.002) | W | 22 | Di-Oxidation | TRUE | TRUE |
| ATCG00270.1 (PSBD/D2) | DIMDDW(1)LR | W | 22 | Tri-Oxidation | TRUE | FALSE |
| ATCG00270.1 (PSBD/D2) | AFNPTQAEETYSM(1)VTANR | M | 247 | Aspartic semialdehyde | FALSE | TRUE |
| ATCG00270.1 (PSBD/D2) | AFNPTQAEETYS(0.023)M(0.78)V(0.191)T(0.006)ANR | M | 247 | Mono-Oxidation | TRUE | TRUE |
| ATCG00270.1 (PSBD/D2) | AYDFV(0.987)S(0.012)Q(0.002)EIR | V | 300 | Mono-Oxidation | TRUE | TRUE |
| ATCG00270.1 (PSBD/D2) | AW(1)MAAQDQPHENLIFPEEVLPR | W | 329 | Kynurenine | TRUE | FALSE |
| ATCG00270.1 (PSBD/D2) | AW(1)MAAQDQPHENLIFPEEVLPR | W | 329 | Hydroxykynurenine | TRUE | FALSE |
| ATCG00270.1 (PSBD/D2) | AW(1)MAAQDQPHENLIFPEEVLPR | W | 329 | Di-Oxidation | TRUE | TRUE |
| ATCG00270.1 (PSBD/D2) | AW(1)MAAQDQPHENLIFPEEVLPR | W | 329 | Tri-Oxidation | TRUE | TRUE |
| ATCG00270.1 (PSBD/D2) | AWM(1)AAQDQPHENLIFPEEVLPR | M | 330 | Aspartic semialdehyde | TRUE | FALSE |
| ATCG00270.1 (PSBD/D2) | M(0.999)AAQDQ(0.001)PHENLIFPEEVLPR | M | 330 | Mono-Oxidation | TRUE | FALSE |
| ATCG00270.1 (PSBD/D2) | AWMAAQ(0.983)DQ(0.983)P(0.034)HENLIFPEEVLPR | Q | 333 | Mono-Oxidation | TRUE | TRUE |
| ATCG00270.1 (PSBD/D2) | AWMAAQ(0.983)DQ(0.983)P(0.034)HENLIFPEEVLPR | Q | 335 | Mono-Oxidation | TRUE | TRUE |
| ATCG00680.1 (PSBB/CP47) | V(1)HTVVLNDPGR | V | 8 | Carbonylation | TRUE | TRUE |
| ATCG00680.1 (PSBB/CP47) | V(0.998)HT(0.002)VVLNDPGR | V | 8 | Mono-Oxidation | TRUE | TRUE |
| ATCG00680.1 (PSBB/CP47) | VH(1)TVVLNDPGR | H | 9 | Aspartic Acid | TRUE | FALSE |
| ATCG00680.1 (PSBB/CP47) | VH(1)TVVLNDPGR | H | 9 | Aspartylurea | TRUE | TRUE |
| ATCG00680.1 (PSBB/CP47) | V(0.992)HT(0.999)V(0.01)VLNDPGR | T | 10 | Mono-Oxidation | TRUE | TRUE |
| ATCG00680.1 (PSBB/CP47) | VHTV(0.001)V(0.003)L(0.981)NDP(0.015)GR(0.001) | L | 13 | Carbonylation | FALSE | TRUE |
| ATCG00680.1 (PSBB/CP47) | V(0.002)HT(0.011)V(0.075)V(0.352)LN(0.863)DP(0.554)GR(0.143) | N | 14 | Mono-Oxidation | TRUE | FALSE |
| ATCG00680.1 (PSBB/CP47) | VHTV(0.001)V(0.027)LN(0.159)DP(0.778)GR(0.035) | P | 16 | Mono-Oxidation | TRUE | FALSE |
| ATCG00680.1 (PSBB/CP47) | Q(0.001)GM(0.999)FVIP(0.596)FM(0.388)T(0.014)R(0.002) | M | 60 | Mono-Oxidation | TRUE | TRUE |
| ATCG00680.1 (PSBB/CP47) | Q(0.006)GM(0.994)FVIP(0.095)FM(0.869)T(0.033)R(0.004) | M | 66 | Mono-Oxidation | FALSE | TRUE |
| ATCG00680.1 (PSBB/CP47) | VSAGLAENQ(0.002)S(0.999)LS(0.999)EAWAK | S | 297 | Mono-Oxidation | TRUE | TRUE |
| ATCG00680.1 (PSBB/CP47) | VSAGLAENQS(0.001)LS(0.999)EAWAK | S | 299 | Mono-Oxidation | TRUE | TRUE |
| ATCG00680.1 (PSBB/CP47) | VSAGLAENQSLSEAW(1)AK | W | 302 | Kynurenine | TRUE | TRUE |
| ATCG00680.1 (PSBB/CP47) | VSAGLAENQSLSEAW(0.976)AK(0.024) | W | 302 | Di-Oxidation | TRUE | FALSE |
| ATCG00680.1 (PSBB/CP47) | DY(0.993)IGNNP(0.005)AK(0.002) | Y | 314 | Di-Oxidation | TRUE | FALSE |
| ATCG00680.1 (PSBB/CP47) | AGS(0.017)M(0.982)DNGDGIAV(0.039)GWLGHP(0.961)V(0.961)FR(0.039) | M | 330 | Mono-Oxidation | TRUE | TRUE |
| ATCG00680.1 (PSBB/CP47) | AGSMDNGDGI(0.999)AV(0.001)GWLGHPVFR | I | 336 | Carbonylation | TRUE | FALSE |
| ATCG00680.1 (PSBB/CP47) | AGSMDNGDGIAV(0.999)GWLGHPVFR | V | 338 | Mono-Oxidation | TRUE | TRUE |
| ATCG00680.1 (PSBB/CP47) | AGSMDNGDGIAVGW(1)LGHPVFR | W | 340 | Di-Oxidation | TRUE | TRUE |
| ATCG00680.1 (PSBB/CP47) | AGS(0.018)M(0.982)DNGDGIAV(0.005)GWLGHP(0.976)V(0.977)FR(0.042) | P | 344 | Mono-Oxidation | TRUE | TRUE |
| ATCG00680.1 (PSBB/CP47) | AGS(0.018)M(0.982)DNGDGIAV(0.005)GWLGHP(0.976)V(0.977)FR(0.042) | V | 345 | Mono-Oxidation | TRUE | TRUE |
| ATCG00680.1 (PSBB/CP47) | AQLGEIFELDR(1) | R | 434 | Mono-Oxidation | TRUE | TRUE |

^1^True, modification present; ^2^False, modification absent; shared modifications are present in both conditions while unique modifications are present only in one condition.

**Table S2.** Shared and unique oxidative modification of monomeric core under H_2_O_2_ and mock

| **Gene (Protein)** | **Modification probabilities** | **Amino acid** |  | **Position** | **Modification** | **Reproducible**  **C + H_2_O_2_** | **Reproducible**  **C + Mock** |
| --- | --- | --- | --- | --- | --- | --- | --- |
| ATCG00020.1 (PSBA/D1) | ESES(1)LWGR(1) | S |  | 12 | Mono-Oxidation | FALSE^1^ | TRUE^2^ |
| ATCG00020.1 (PSBA/D1) | ESESLW(0.991)GR(0.009) | W |  | 14 | Di-Oxidation | TRUE | TRUE |
| ATCG00020.1 (PSBA/D1) | ESES(1)LWGR(1) | R |  | 16 | Mono-Oxidation | TRUE | TRUE |
| ATCG00020.1 (PSBA/D1) | FCN(1)WITSTENR | N |  | 19 | Mono-Oxidation | TRUE | TRUE |
| ATCG00020.1 (PSBA/D1) | FCN(0.003)WIT(0.925)S(0.923)T(0.148)EN(0.001)R | T |  | 22 | Mono-Oxidation | TRUE | TRUE |
| ATCG00020.1 (PSBA/D1) | FCN(0.003)WIT(0.925)S(0.923)T(0.148)EN(0.001)R | S |  | 23 | Mono-Oxidation | FALSE | TRUE |
| ATCG00020.1 (PSBA/D1) | EW(1)ELSFR | W |  | 131 | Kynurenine | TRUE | FALSE |
| ATCG00020.1 (PSBA/D1) | EW(1)ELSFR | W |  | 131 | Di-Oxidation | FALSE | TRUE |
| ATCG00020.1 (PSBA/D1) | EWELS(0.999)FR(0.001) | S |  | 134 | Mono-Oxidation | TRUE | FALSE |
| ATCG00020.1 (PSBA/D1) | ET(0.001)T(0.067)EN(0.15)ES(0.497)AN(0.837)EGYR(0.448) | N |  | 234 | Mono-Oxidation | TRUE | TRUE |
| ATCG00020.1 (PSBA/D1) | ETTENESAN(0.003)EGYR(0.997) | R |  | 238 | Mono-Oxidation | FALSE | TRUE |
| ATCG00020.1 (PSBA/D1) | VIN(1)T(1)WADIINR | N |  | 315 | Mono-Oxidation | TRUE | TRUE |
| ATCG00020.1 (PSBA/D1) | VIN(1)T(1)WADIINR | T |  | 316 | Mono-Oxidation | TRUE | TRUE |
| ATCG00020.1 (PSBA/D1) | VINTW(1)ADIINR | W |  | 317 | Kynurenine | TRUE | TRUE |
| ATCG00020.1 (PSBA/D1) | VINTW(1)ADIINR | W |  | 317 | Oxolactone | TRUE | FALSE |
| ATCG00020.1 (PSBA/D1) | VINTW(1)ADIINR | W |  | 317 | Di-Oxidation | TRUE | TRUE |
| ATCG00020.1 (PSBA/D1) | VINTW(1)ADIINR | W |  | 317 | Tri-Oxidation | TRUE | TRUE |
| ATCG00020.1 (PSBA/D1) | V(0.012)I(0.012)NTWADI(0.877)I(0.098)NR(0.002) | I |  | 320 | Carbonylation | TRUE | FALSE |
| ATCG00020.1 (PSBA/D1) | VIN(0.003)T(0.003)WADIIN(0.997)R(0.997) | N |  | 322 | Mono-Oxidation | TRUE | TRUE |
| ATCG00020.1 (PSBA/D1) | VIN(0.003)T(0.003)WADIIN(0.997)R(0.997) | R |  | 323 | Mono-Oxidation | TRUE | TRUE |
| ATCG00020.1 (PSBA/D1) | ANLGM(1)EVMHER | M |  | 328 | Mono-Oxidation | TRUE | TRUE |
| ATCG00020.1 (PSBA/D1) | AN(0.001)LGMEV(0.995)MHER(0.004) | V |  | 330 | Mono-Oxidation | TRUE | TRUE |
| ATCG00020.1 (PSBA/D1) | ANLGM(0.98)EV(0.026)M(0.993)HER(0.001) | M |  | 331 | Mono-Oxidation | TRUE | TRUE |
| ATCG00270.1 (PSBD/D2) | DLFDIM(1)DDWLR | M |  | 19 | Mono-Oxidation | TRUE | FALSE |
| ATCG00270.1 (PSBD/D2) | DIMDDW(0.998)LR(0.002) | W |  | 22 | Di-Oxidation | FALSE | TRUE |
| ATCG00270.1 (PSBD/D2) | DIMDDW(1)LR | W |  | 22 | Tri-Oxidation | TRUE | FALSE |
| ATCG00270.1 (PSBD/D2) | AFNPTQAEETYS(0.023)M(0.78)V(0.191)T(0.006)ANR | M |  | 247 | Mono-Oxidation | FALSE | TRUE |
| ATCG00270.1 (PSBD/D2) | AYDFV(0.987)S(0.012)Q(0.002)EIR | V |  | 300 | Mono-Oxidation | TRUE | TRUE |
| ATCG00270.1 (PSBD/D2) | AW(1)MAAQDQPHENLIFPEEVLPR | W |  | 329 | Di-Oxidation | TRUE | TRUE |
| ATCG00270.1 (PSBD/D2) | AW(1)MAAQDQPHENLIFPEEVLPR | W |  | 329 | Tri-Oxidation | TRUE | TRUE |
| ATCG00270.1 (PSBD/D2) | M(0.999)AAQDQ(0.001)PHENLIFPEEVLPR | M |  | 330 | Mono-Oxidation | FALSE | TRUE |
| ATCG00270.1 (PSBD/D2) | AWMAAQ(0.983)DQ(0.983)P(0.034)HENLIFPEEVLPR | Q |  | 333 | Mono-Oxidation | TRUE | TRUE |
| ATCG00270.1 (PSBD/D2) | AWMAAQ(0.983)DQ(0.983)P(0.034)HENLIFPEEVLPR | Q |  | 335 | Mono-Oxidation | TRUE | TRUE |
| ATCG00680.1 (PSBB/CP47) | V(0.05)HT(0.125)V(0.873)V(0.846)LN(0.07)DP(0.036)GR | V |  | 11 | Mono-Oxidation | FALSE | TRUE |
| ATCG00680.1 (PSBB/CP47) | VHTVVLNDP(0.996)GR(0.004) | P |  | 16 | Di-Oxidation | FALSE | TRUE |
| ATCG00680.1 (PSBB/CP47) | ELAVFDPSDPVLDPMW(1)R | W |  | 56 | Tri-Oxidation | TRUE | FALSE |
| ATCG00680.1 (PSBB/CP47) | Q(0.001)GM(0.999)FVIP(0.596)FM(0.388)T(0.014)R(0.002) | M |  | 60 | Mono-Oxidation | TRUE | TRUE |
| ATCG00680.1 (PSBB/CP47) | VSAGLAENQ(0.002)S(0.999)LS(0.999)EAWAK | S |  | 297 | Mono-Oxidation | TRUE | TRUE |
| ATCG00680.1 (PSBB/CP47) | VSAGLAENQS(0.001)LS(0.999)EAWAK | S |  | 299 | Mono-Oxidation | TRUE | TRUE |
| ATCG00680.1 (PSBB/CP47) | VSAGLAENQSLSEAW(0.976)AK(0.024) | W |  | 302 | Di-Oxidation | TRUE | FALSE |
| ATCG00680.1 (PSBB/CP47) | AGSMDNGDGI(0.999)AV(0.001)GWLGHPVFR | I |  | 336 | Carbonylation | FALSE | TRUE |
| ATCG00680.1 (PSBB/CP47) | AGSMDNGDGIAV(0.999)GWLGHPVFR | V |  | 338 | Mono-Oxidation | TRUE | TRUE |
| ATCG00680.1 (PSBB/CP47) | AGSMDNGDGIAVGW(1)LGHPVFR | W |  | 340 | Di-Oxidation | TRUE | TRUE |
| ATCG00680.1 (PSBB/CP47) | AQLGEIFELDR(1) | R |  | 434 | Mono-Oxidation | TRUE | TRUE |

^1^False, modification absent; ^2^True, modification present; shared modifications are present in both conditions while unique modifications are present only in one condition.

**Table S3.** Core phosphoproteins, phosphopeptides, and phosphosites of *Arabidopsis* PSII

| **Gene ID (Protein)** | **Phospho (S/T) probabilities** | **Position within proteins** | **Amino acid** |
| --- | --- | --- | --- |
| AT1G79040.1 (PsbR) | TDKPFGINGS(1)MDLR | 58 | S |
| AT4G05180.1 (PsbQ-2) | FYIQPLS(0.966)PT(0.034)EAAAR | 125 | S |
| ATCG00020.1 (PsbA/D1) | T(1)AILERR | 2 | T |
| ATCG00270.1 (PsbD/D2) | T(1)IALGK | 2 | T |
| ATCG00280.1 (PsbC/CP43) | GIDRDFEPVLS(0.5)MT(0.5)PLN | 468 | S |
| ATCG00280.1 (PsbC/CP43) | GIDRDFEPVLS(0.5)MT(0.5)PLN | 470 | T |
| ATCG00560.1 (PsbL) | T(1)QSNPNEQSVELNR | 2 | T |
| ATCG00570.1 (PsbF) | T(0.983)IDRT(0.017)YPIFTVR | 2 | T |
| ATCG00710.1 (PsbH) | AT(1)QT(0.999)VEDSSR | 3 | T |
| ATCG00710.1 (PsbH) | AT(1)QT(0.999)VEDSSR | 5 | T |
